# Structural and functional analysis of cancer-associated missense variants in the retinoblastoma protein (Rb) pocket domain

**DOI:** 10.1101/2024.09.22.614354

**Authors:** Anthony Castro, Alfredo Ruiz-Rivera, Chad C. Moorman, Emma R. Wolf-Saxon, Hailey N. Mims, Vanessa I. Vasquez-Meza, Matthew A. Rangel, Marcos M. Loera, Ian Bond, Seth B. Buchanan, Estela Villarreal, Sarvind Tripathi, Seth M. Rubin, Jason R. Burke

**Affiliations:** Department of Chemistry and Biochemistry, California State University, San Bernardino, CA 92407, USA; Department of Chemistry and Biochemistry, University of California, Santa Cruz, CA 95064, USA

**Keywords:** Cancer missense mutation, cell cycle, transcription regulation, retinoblastoma protein, RB, E2F, LxCxE

## Abstract

The retinoblastoma tumor suppressor (Rb) is a multifunctional protein that primarily regulates the cell cycle but also has roles in cellular differentiation, DNA damage response and apoptosis. The loss of Rb is a key event in the development or progression of many cancers. Essential functions of Rb occur through its pocket domain, which is necessary for regulating binding interactions with E2F transcription factors and transcription repressors that bind via an LxCxE motif. The pocket domain is the most highly-conserved region of the multidomain protein, as well as the most frequent site of mutations. To understand what effects cancer missense mutations have on Rb’s pocket domain, we used fluorescence polarization and differential scanning fluorimetry to quantify changes, caused by 75 cancer-associated missense variants, to E2F transactivation domain (E2F^TD^) binding, LxCxE binding, and changes to the thermostability of the protein. We find that 43% of the missense variants we tested reduce Rb-E2F^TD^ binding. Many of these variants are not located at the E2F binding site, yet they destabilize the fold of the protein and show temperature-sensitive binding effects. We also find that 21% of tested mutations reduce LxCxE binding, and several mutations selectively disrupt either E2F^TD^ or LxCxE binding. Protein X-ray crystallography of four missense variants reveals how different mutations destabilize the protein fold and inhibit E2F^TD^ or LxCxE binding. Taken together, this work provides the first understanding of the multiple ways through which stability, structure and function of Rb’s pocket domain is altered by a large number of missense mutations seen in cancer.

## INTRODUCTION

The retinoblastoma protein (Rb) is a multifunctional, chromatin associated tumor suppressor protein. Functional Rb is critical for preventing the initiation of retinoblastomas, small cell lung cancers and osteosarcoma, and Rb loss correlates with cancer progression of most other common human cancers (**Burkhart 2008**). Recent reviews highlight the numerous context-dependent, noncanonical and enigmatic roles of Rb and its deletion in cancers (**Dyson 2016, Dick 2018, Cobrinik 2024**). However, Rb’s most critical role is as a key regulator of the cell cycle (**Rubin 2020**). One component of this regulation occurs late in G_1_ phase, when hyperphosphorylation of Rb by cyclin-CDKs disrupts repressive complexes between Rb and E2F, resulting in the activation of S-phase gene transcription (**Rubin 2020).** Phosphorylation causes structural changes to Rb that strongly inhibit E2F transactivation domain binding (E2F^TD^) (**Rubin 2012, Burke 2014)**. Underscoring the importance of these inactivation mechanisms, Rb hyperphosphorylation is a common consequence of cancerous Rb pathway mutations **(Sherr 2016**). In certain contexts, cancer-associated missense mutations may be similar to the effects of persistent hyperphosphorylation of Rb, as they may enable retention of select functions of folded Rb while disrupting specific protein-protein interactions. This idea has been tested for some mutations which weaken Rb-E2F^TD^ binding interactions *in vitro* (**Shan 1996, Otterson 1997**), and *in vivo* (**Sellers 1998**). However, there is currently little known about the biochemical consequences of a majority of the observed cancer-associated missense mutations and to what extent these affect Rb-E2F and other Rb complexes.

Genome sequencing efforts have revealed detailed landscapes of somatic mutations in cancers (**Cancer Genome Atlas Research Network 2013**). For a majority of these, clinical, cellular, biochemical and structural outcomes remain unclear. Within the Rb protein, hundreds of cancer-associated missense mutations span multiple structured domains comprising conserved protein-protein interaction sites that regulate distinct functions. (**Dick 2013, Topacio 2019**). Among these are mutations which potentially affect the structured LxCxE-binding cleft, which is responsible for recruiting co-repressive complexes to silence E2F promoters (**Dick 2013**). Mutation studies specifically targeting the LxCxE binding site have parsed some of Rb’s context-dependent tumor suppressive roles onto distinct regions of the protein. For example, interactions at the LxCxE binding site do not affect cell cycle arrest, differentiation or repression of E2F targets, but instead prevent entry into senescence (**Talluri 2010, Thwaites 2017**). However, in the context of genotoxic stress caused by DNA damage, mutations specifically targeting the LxCxE binding site do compromise repression of E2F targets and cell cycle control (**Bourgo 2011, Talluri 2013, Volmer 2014**). Without genotoxic stress, a combination of mutations at multiple protein-protein interaction sites is needed to maximally impair Rb mediated cell cycle control (**Thwaites 2017**). The proteins that interact directly with Rb via LxCxE motifs include tumor suppressors ARID4A and KDM5A, as well as EID1 and others (**Putta 2022**). Multiple oncogenic viruses have convergently evolved to target Rb via the LxCxE interface, suggesting an importance of the tumor suppressive functions at the site (**Felsani 2006**). One well-studied example is the E7 protein made by the Human Papilloma Virus (**Tomita 2020**). Nevertheless, on the molecular and structural levels, it is still not entirely clear how Rb coordinates its multiple roles, and which functions are lost that drive the initiation and evolution and specific cancers (**Dyson 2016**). The M704V and R661W cancer mutations have been shown to disrupt LxCxE-mediated protein binding; however, it has been unclear whether other cancer mutations affect binding interactions as this site as well (**Otterson 1997**, **Henley 2010**).

Studies of specific cancer-associated Rb missense mutations have provided some insights into their consequences. The most well studied example, R661W, is a low-penetrance mutation in familial retinoblastoma that contributes partially to cell cycle arrest in retinoblastoma, is sufficient to initiate tumorigenesis, and causes cell cycle defects in embryos (**Cobrinik 2006, Sun 2006, Sun 2011**). Additionally, R661W, which alters both E2F and LxCxE binding, exhibits temperature sensitivity in a yeast growth assay, such that LxCxE-binding is reduced at 30°C and 37°C when compared with 25°C (**Otterson 1997**, **Otterson 1999**). Temperature dependence is also a characteristic of some cancer-associated missense mutations in p53, where variants that destabilize p53 also disrupt binding to DNA, enhance degradation and are drug targets for intervention by stabilizing compounds (**Fallatah 2023**). Beyond R661W, however, for Rb there has not been evidence of the consequences that cancer missense mutations have on thermostability and related changes to important protein-protein interactions (**Otterson 1999**).

The goal of this work is to use biochemical approaches to characterize the effects of a large set of the most common cancer-associated missense variants that occur to the pocket domain of the retinoblastoma tumor suppressor protein. By creating a large data set of binding and stability measurements of Rb missense variants, we seek to demystify predictive qualities about how these variants may differently affect Rb’s functions. The pocket domain was selected because it is the most frequent target of cancer mutations, it contains multiple functional binding interfaces, and it comprises a folded domain that is amenable to study through biochemical means. While a handful of cancer missense mutations to the pocket domain have been studied for E2F and LxCxE binding defects, there is currently little known about the biochemical consequences of the vast majority of mutations associated with cancer (**Shan 1996, Otterson 1997, Henley 2010**). Here, we profile 75 Rb missense variants for changes to key binding interactions. We use E7^LXCXE^ in order to understand effects that may occur to structurally-similar, endogenous LxCxE-mediated binding interactions (**Putta 2022**). The E7 oncoprotein from HPV has provided a structured example of binding at Rb’s LxCxE site, and has been used extensively as a tool to study and characterize binding at this site (**Lee 1998, Chemes 2010, Pye 2016**). In addition, we measure missense variant-induced changes to E2F1^TD^ binding, which may serve as an indicator for binding changes to the homologous activating E2Fs. Thermostability changes and temperature sensitive binding effects are also measured, and together with protein X-ray crystallography, these experiments provide a comprehensive understanding of the biochemical consequences of cancer missense mutations to Rb’s pocket domain, as well as trends that define them.

## RESULTS

### Expression of Rb Pocket Domain Missense Variants in *e coli*

For this study, 112 missense variants in the Rb pocket domain (Rb^P^) were identified from two pan cancer genome somatic mutation databases, COSMIC (cancer.sanger.ac.uk) **(Tate 2019**), and cBioPortal (cbioportal.org) (**Cerami 2012)**. After cloning and expression attempts, we found that 37 missense variants could not be expressed from *e coli*, indicating that protein misfolding or severe instability may be caused by these mutations (**Figure 1**). The remaining 75 variants were successfully expressed and purified to homogeneity from *e coli*. The studied variants span all regions of the pocket domain, which includes subdomain A, subdomain B and the unstructured 65-amino acid “pocket loop” (Rb^PL^) (**Figure 1**).

**Figure 1.**
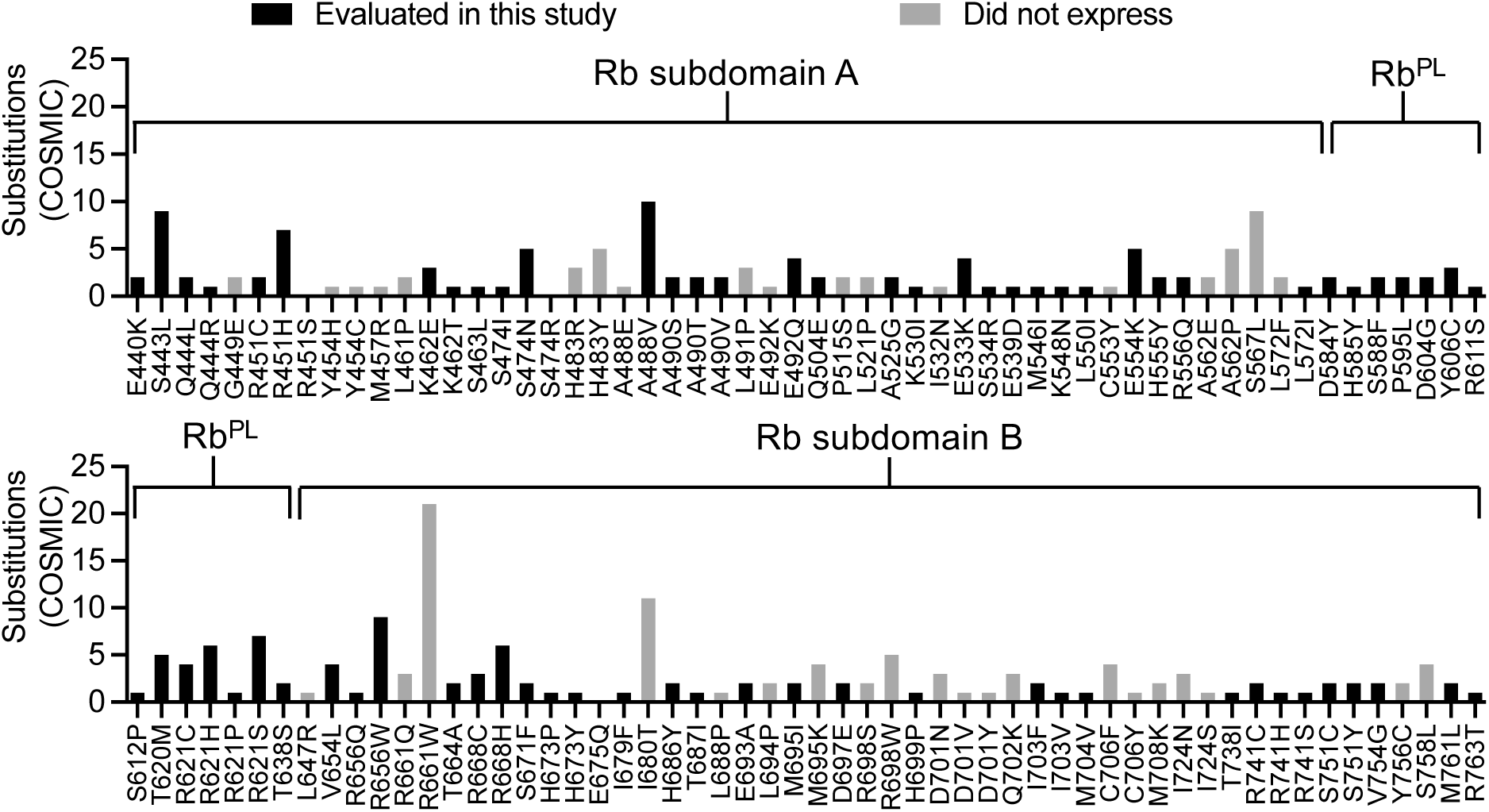
Missense variants of human Rb used or attempted in this study. Each missense variant was independently cloned, expressed and evaluated as a GST-Rb^P^ fusion protein from *e coli*. Missense variants that failed to produce soluble protein from this system are shown as grey bars (n=37). The variants that did express in the host system and underwent further biochemical characterizations are shown as black bars (n=75). For each variant, substitutions (numbers of incidents in patients) are taken from the COSMIC somatic cancer mutation dataset and are shown on the y-axis. Variants that have neither grey nor black bars are taken from the cBioPortal data set; these were successfully expressed and characterized in this study.

### Disruptions to Protein-Protein Interactions are a Common Consequence of Cancer-associated Rb^P^ Missense Variants

Changes to binding affinities caused by missense variants were measured by fluorescence polarization (FP). In this FP assay, a peptide of the E2F1 transactivation domain that is N-labeled with tetramethyl rhodamine dye (TMR-E2F1^TD^), is measured in the presence of unlabeled Rb^P^ (**Pye 2016**). When the experiment is conducted at 25°C, the measured equilibrium dissociation constant (K_D_) for E2F1^TD^ binding to wild type Rb^P^ is 14 ± 3 nM (**Table 1**, **Figure 2A**). This value is similar to the K_D_ value measured for wild type Rb^P^-E2F1^TD^ by isothermal titration calorimetry (ITC) (**Burke 2010**). To examine Rb^P^ missense variants in this manner, we first considered that mutations to proteins tend to be destabilizing and can cause irreversible unfolding. In a binding assay, a tendency for unfolding can have the consequence of reducing the known concentration of active protein used for K_D_ measurements, necessitating the reporting of apparent K_D_ values (K_D (app)_), as opposed to K_D_ values (**Altschuler 2013, Jarmoskaite 2020**). In its wild type form, the construct Rb^P^ is metastable when purified *in vitro*, and the protein is prone to temperature dependent irreversible unfolding transitions (**Chemes 2013**). Taking this into consideration, we here report apparent equilibrium dissociation constant values (K_D (app)_) for each Rb^P^ variant. Through these measurements, we are able to compare changes to the binding capabilities of Rb^P^ missense variants. This type of comparison is particularly important for evaluating cancer-associated mutations at physiological temperature (37°C), where effects of the variants are most relevant.

**Figure 2.**
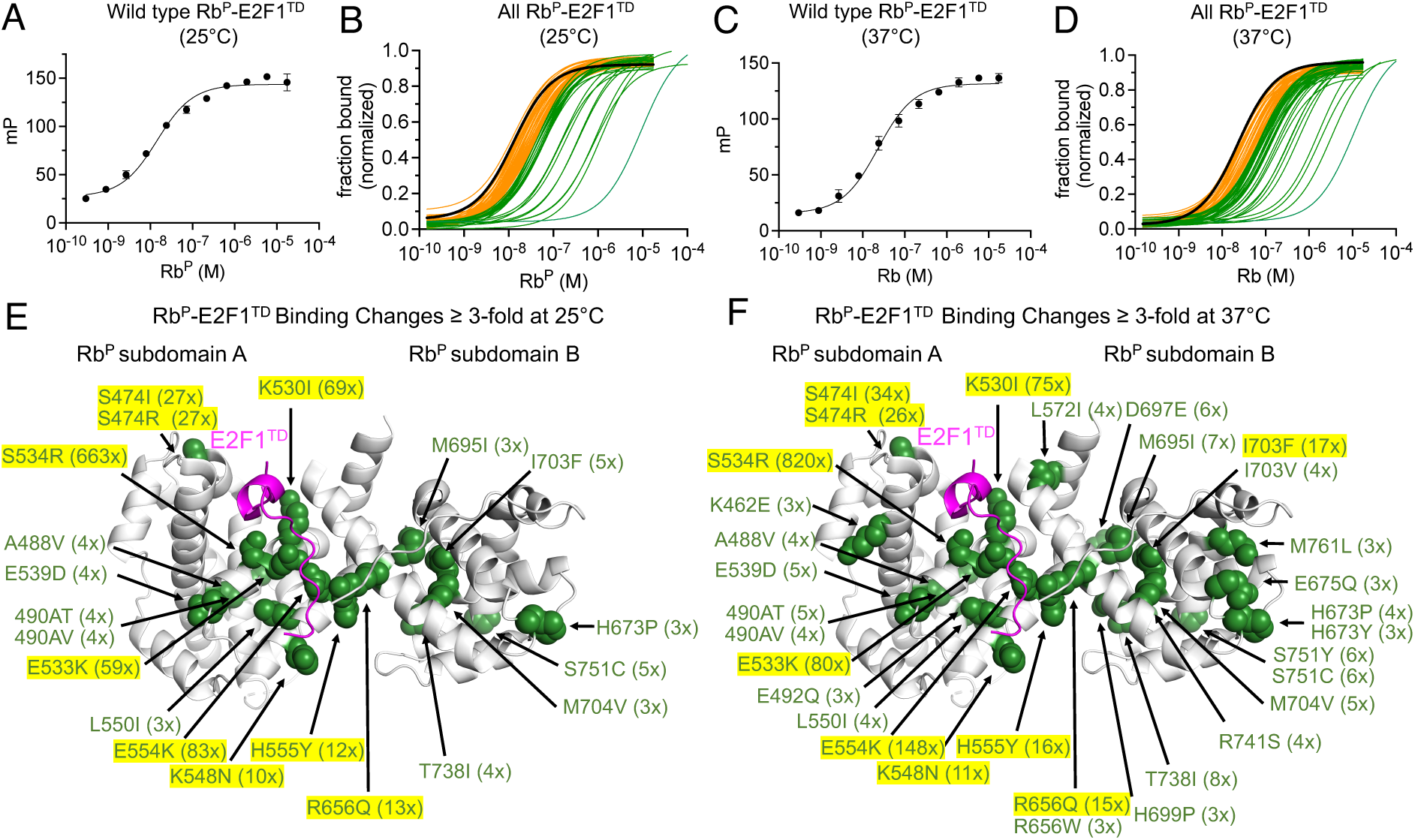
Fluorescence polarization binding experiments for Rb^P^-E2F1^TD^ at 25°C and 37°C. **A**) The saturating binding curve for TMR-E2F1^TD^ binding to wild type Rb^P^ at 25°C. Error bars are standard deviations of four replicates. **B**) Normalized binding curves for Rb^P^-E2F1^TD^ at 25°C for: wild type Rb^P^ (black, n=1); Rb^P^ variants which show less than 3-fold increases to K_D (app)_ relative to wild type (orange, n=55); and Rb^P^ variants which show 3-fold or greater increases to K_D (app)_ relative to wild type (green, n=20). **C**) The saturating binding curve for TMR-E2F1^TD^ binding to wild type Rb^P^ at 37°C. Error bars are standard deviations of four replicates. **D**) Normalized binding curves for Rb^P^-E2F1^TD^ at 37°C for: wild type Rb (black, n=1); Rb^P^ variants which show less than 3-fold increases to K_D (app)_ relative to wild type (orange, n=43); and Rb^P^ variants which show 3-fold or greater increases to K_D (app)_ (green, n=32). The structural positions of the missense variants (green, space-filling) with 3-fold or greater increases to K_D (app)_ are mapped onto the structure of Rb^P^-E2F1^TD^ (PDB: 1O9K) (**Xiao 2003**) for 25°C binding experiments (**E**) and 37°C binding experiments (**F**). Fold increases in K_D (app)_ relative to wild type are shown in parentheses next to each mutation name, and those with greater than 10-fold effects are highlighted.

**Table 1.** K_D (app)_ and T_m_ data for all Rb^P^ variants and wild type (*denotes K_D_ value). Reported errors in the K_D_ or K_D (app)_ from FP measurements are curve-fitting errors.

| Rb<sup>P</sup> variant | Rb<sup>P</sup> - E2F1<sup>TD</sup> $K_D$ (app) (nM) | | Rb<sup>P</sup> - E7<sup>LxCxE</sup> $K_D$ (app) (nM) | | $T_m$ (°C) |
| --- | --- | --- | --- | --- | --- |
|  | 25°C | 37°C | 25°C | 37°C |  |
| **Wild type** | **14 ± 3\*** | **22 ± 4** | **5 ± 1\*** | **12 ± 1** | **46.9 ± 0.5** |
| 1 E440K | 12 ± 3 | 27 ± 4 | 5 ± 1 | 13 ± 1 | 44.2 ± 0.7 |
| 2 S443L | 31 ± 6 | 55 ± 11 | 9 ± 1 | 21 ± 2 | 45.4 ± 0.3 |
| 3 Q444L | 14 ± 3 | 30 ± 7 | 4 ± 1 | 11 ± 2 | 46.0 ± 0.8 |
| 4 Q444R | 15 ± 3 | 25 ± 4 | 7 ± 2 | 17 ± 4 | 47.3 ± 0.6 |
| 5 R451C | 33 ± 7 | 51 ± 9 | 7 ± 2 | 19 ± 4 | 46.4 ± 1.1 |
| 6 R451H | 29 ± 6 | 56 ± 13 | 9 ± 2 | 22 ± 4 | 44.4 ± 0.7 |
| 7 R451S | 26 ± 5 | 44 ± 9 | 9 ± 2 | 17 ± 2 | 45.5 ± 0.4 |
| 8 K462E | 27 ± 4 | 67 ± 17 | 11 ± 2 | 32 ± 9 | 41.9 ± 0.2 |
| 9 K462T | 30 ± 6 | 51 ± 10 | 8 ± 1 | 16 ± 3 | 47.2 ± 0.0 |
| 10 S463L | 26 ± 5 | 41 ± 7 | 10 ± 7 | 17 ± 12 | 48.0 ± 0.8 |
| 11 S474I | 364 ± 53 | 775 ± 148 | 7 ± 1 | 18 ± 2 | 43.7 ± 0.1 |
| 12 S474N | 19 ± 5 | 26 ± 7 | 8 ± 3 | 13 ± 4 | 45.7 ± 0.3 |
| 13 S474R | 365 ± 37 | 580 ± 63 | 5 ± 1 | 15 ± 3 | 44.4 ± 0.1 |
| 14 A488V | 53 ± 11 | 78 ± 17 | 9 ± 1 | 19 ± 2 | 42.8 ± 0.3 |
| 15 A490S | 17 ± 3 | 29 ± 5 | 7 ± 1 | 15 ± 1 | 46.1 ± 0.3 |
| 16 A490T | 59 ± 13 | 115 ± 32 | 13 ± 2 | 33 ± 4 | 42.2 ± 0.9 |
| 17 A490V | 48 ± 11 | 87 ± 20 | 11 ± 1 | 33 ± 3 | 41.8 ± 0.3 |
| 18 E492Q | 22 ± 5 | 73 ± 18 | 14 ± 3 | 52 ± 12 | 41.5 ± 0.6 |
| 19 Q504E | 18 ± 5 | 33 ± 9 | 5 ± 1 | 11 ± 2 | 46.9 ± 0.3 |
| 20 A525G | 36 ± 7 | 65 ± 12 | 12 ± 4 | 52 ± 13 | 47.7 ± 0.4 |
| 21 K530I | 945 ± 296 | 1680 ± 475 | 8 ± 3 | 20 ± 6 | 46.9 ± 0.2 |
| 22 E533K | 810 ± 153 | 1805 ± 277 | 6 ± 1 | 14 ± 2 | 40.5 ± 0.1 |
| 23 S534R | 9278 ± 2438 | 18050 ± 3072 | 4 ± 2 | 11 ± 3 | 45.6 ± 0.3 |
| 24 E539D | 55 ± 11 | 104 ± 17 | 7 ± 1 | 19 ± 2 | 42.0 ± 0.2 |
| 25 M546I | 13 ± 3 | 21 ± 4 | 7 ± 1 | 13 ± 1 | 47.2 ± 0.2 |
| 26 K548N | 133 ± 39 | 178 ± 56 | 6 ± 1 | 15 ± 3 | 46.5 ± 0.1 |
| 27 L550I | 41 ± 6 | 80 ± 10 | 3 ± 1 | 7 ± 1 | 43.2 ± 0.5 |
| 28 E554K | 1167 ± 98 | 3266 ± 722 | 12 ± 3 | 48 ± 9 | 39.9 ± 0.2 |
| 29 H555Y | 158 ± 28 | 366 ± 72 | 8 ± 2 | 32 ± 6 | 44.4 ± 0.1 |
| 30 R556Q | 22 ± 5 | 48 ± 12 | 5 ± 1 | 13 ± 2 | 46.7 ± 0.5 |
| 31 L572I | 37 ± 12 | 87 ± 30 | 4 ± 2 | 15 ± 8 | 47.2 ± 0.7 |
| 32 D584Y | 33 ± 7 | 51 ± 12 | 12 ± 2 | 20 ± 4 | 46.2 ± 0.1 |
| 33 H585Y | 35 ± 8 | 92 ± 40 | 6 ± 1 | 18 ± 2 | 47.4 ± 0.8 |
| 34 S588F | 36 ± 8 | 51 ± 12 | 5 ± 1 | 16 ± 1 | 46.0 ± 0.1 |
| 35 P595L | 46 ± 9 | 83 ± 12 | 14 ± 3 | 32 ± 6 | 46.8 ± 0.2 |
| 36 D604G | 27 ± 5 | 50 ± 11 | 13 ± 5 | 26 ± 9 | 46.2 ± 0.7 |

| Rb<sup>P</sup> variant | Rb<sup>P</sup>- E2F1<sup>TD</sup> K<sub>D (app)</sub> (nM) |  | Rb<sup>P</sup>- E7<sup>LxCxE</sup> K<sub>D (app)</sub> (nM) |  | T<sub>m</sub> (°C) |
| --- | --- | --- | --- | --- | --- |
|  | 25°C | 37°C | 25°C | 37°C |  |
| 37 Y606C | 15 ± 3 | 27 ± 4 | 7 ± 4 | 23 ± 11 | 45.4 ± 0.3 |
| 38 R611S | 15 ± 4 | 30 ± 11 | 5 ± 1 | 15 ± 5 | 46.7 ± 0.1 |
| 39 S612P | 32 ± 6 | 49 ± 11 | 7 ± 2 | 17 ± 3 | 45.5 ± 0.1 |
| 40 T620M | 24 ± 8 | 59 ± 16 | 6 ± 2 | 15 ± 5 | 46.1 ± 1.1 |
| 41 R621C | 22 ± 4 | 43 ± 7 | 12 ± 3 | 30 ± 5 | 45.8 ± 0.0 |
| 42 R621H | 30 ± 5 | 48 ± 8 | 10 ± 3 | 18 ± 4 | 46.9 ± 0.1 |
| 43 R621P | 40 ± 8 | 83 ± 20 | 12 ± 2 | 32 ± 6 | 45.9 ± 0.6 |
| 44 R621S | 21 ± 5 | 38 ± 9 | 6 ± 1 | 17 ± 3 | 45.9 ± 0.4 |
| 45 T638S | 14 ± 2 | 38 ± 9 | 7 ± 1 | 14 ± 3 | 46.5 ± 0.2 |
| 46 S648T | 33 ± 9 | 58 ± 14 | 6 ± 1 | 14 ± 3 | 46.6 ± 0.2 |
| 47 V654L | 18 ± 3 | 30 ± 4 | 11 ± 2 | 21 ± 2 | 48.2 ± 0.1 |
| 48 R656Q | 183 ± 36 | 325 ± 75 | 6 ± 1 | 14 ± 1 | 46.5 ± 0.1 |
| 49 R656W | 39 ± 7 | 77 ± 11 | 8 ± 2 | 19 ± 3 | 43.1 ± 0.2 |
| 50 T664A | 15 ± 2 | 24 ± 4 | 8 ± 4 | 16 ± 5 | 46.6 ± 0.1 |
| 51 R668C | 12 ± 4 | 25 ± 10 | 3 ± 1 | 16 ± 3 | 44.9 ± 0.6 |
| 52 R668H | 13 ± 2 | 22 ± 4 | 7 ± 3 | 28 ± 7 | 44.2 ± 0.1 |
| 53 S671F | 24 ± 5 | 46 ± 11 | 9 ± 1 | 19 ± 2 | 46.1 ± 0.2 |
| 54 H673P | 42 ± 6 | 98 ± 15 | 24 ± 2 | 83 ± 6 | 43.3 ± 0.8 |
| 55 H673Y | 26 ± 5 | 66 ± 13 | 10 ± 1 | 29 ± 3 | 45.2 ± 0.0 |
| 56 E675Q | 35 ± 8 | 71 ± 21 | 20 ± 9 | 46 ± 18 | 46.4 ± 0.1 |
| 57 I679F | 17 ± 4 | 33 ± 10 | 11 ± 5 | 20 ± 10 | 45.9 ± 0.1 |
| 58 H686Y | 20 ± 4 | 35 ± 7 | 5 ± 1 | 14 ± 2 | 46.6 ± 0.5 |
| 59 T687I | 20 ± 4 | 40 ± 6 | 13 ± 1 | 29 ± 2 | 48.9 ± 0.3 |
| 60 E693A | 42 ± 6 | 59 ± 18 | 7 ± 1 | 24 ± 3 | 43.1 ± 0.2 |
| 61 M695I | 43 ± 10 | 165 ± 33 | 11 ± 6 | 93 ± 44 | 43.8 ± 0.2 |
| 62 D697E | 29 ± 6 | 129 ± 35 | 9 ± 2 | 46 ± 8 | 42.8 ± 0.2 |
| 63 H699P | 35 ± 6 | 68 ± 14 | 6 ± 1 | 14 ± 2 | 43.6 ± 0.2 |
| 64 I703F | 63 ± 12 | 396 ± 12 | 16 ± 2 | 275 ± 26 | 40.3 ± 0.4 |
| 65 I703V | 20 ± 4 | 99 ± 22 | 21 ± 11 | 53 ± 26 | 43.7 ± 0.2 |
| 66 M704V | 41 ± 8 | 129 ± 37 | 18 ± 4 | 51 ± 16 | 41.8 ± 0.7 |
| 67 T738I | 60 ± 11 | 179 ± 34 | 18 ± 2 | 151 ± 24 | 41.6 ± 0.1 |
| 68 R741C | 23 ± 3 | 64 ± 12 | 14 ± 3 | 23 ± 2 | 44.3 ± 0.2 |
| 69 R741H | 21 ± 4 | 41 ± 7 | 18 ± 6 | 21 ± 4 | 45.8 ± 0.1 |
| 70 R741S | 32 ± 7 | 84 ± 23 | 16 ± 1 | 46 ± 5 | 43.7 ± 0.4 |
| 71 S751C | 66 ± 13 | 137 ± 32 | 44 ± 7 | 98 ± 19 | 46.0 ± 0.4 |
| 72 S751Y | 33 ± 5 | 127 ± 25 | 127 ± 8 | 420 ± 38 | 41.5 ± 0.2 |
| 73 V754G | 20 ± 4 | 34 ± 7 | 7 ± 2 | 15 ± 2 | 44.6 ± 0.3 |
| 74 M761L | 32 ± 7 | 68 ± 13 | 1.3 ± 0.4 | 5 ± 1 | 46.4 ± 0.3 |
| 75 R763T | 37 ± 11 | 59 ± 18 | 21 ± 6 | 35 ± 7 | 45.3 ± 0.3 |

When the Rb^P^-E2F1^TD^ binding experiment is conducted using 75 missense variants of Rb^P^ at 25°C, we observe that 27% of the variants tested (n=20) have 3-fold, or more, weaker binding to E2F1^TD^ relative to wild type (**Table 1**, **Figure 2B, Supplemental Figure 1).** Although *in vitro* binding assays are most often conducted under standard biochemical temperature at 25°C, the effects of certain cancer-associated mutations may be observable only at physiological temperature (37°C). To address this possibility, the experiments were conducted again at 37°C. For wild type Rb^P^ binding to E2F1^TD^, the measured apparent equilibrium dissociation constant (K_D (app)_) at physiological temperature is 22 ± 4 nM (**Table 1**, **Figure 2C**). When the same experiment is conducted using 75 Rb^P^ missense variants, we observe that 43% of the variants tested (n=32) cause 3-fold, or more, weaker E2F1^TD^ binding relative to wild type (**Table 1**, **Figure 2D, Supplemental Figure 1**). These data sets directly overlap, such that the variants with 3-fold or greater effects at 25°C also have 3-fold or greater effects at 37°C. The large percentage of tested mutations that are found to weaken Rb^P^ binding to E2F1^TD^ reveals how broadly susceptible this important binding interaction is to disruptions caused by cancer-associated missense mutations in Rb^P^.

Trends emerge when the positions of missense variants are mapped onto the structure of Rb’s pocket domain. At 25°C, the majority of variants that strongly reduce E2F1^TD^ binding are located at the E2F^TD^ binding cleft; these include S534R (820-fold), K530I (63-fold); E533K (59-fold); E544K (34-fold); R656Q (13-fold); H555Y (12-fold); and K548N (10-fold) (**Figure 2E**). In the X-ray structures of Rb^P^-E2F1^TD^ and Rb^P^-E2F2^TD^, each of the wild type forms of these 7 amino acids forms binding interactions with E2F^TD^; however, at that time these amino acids were not known as missense mutation sites in cancers (**Lee 2002, Xiao 2003**). Two variants show greater than 10-fold reductions in binding to E2F1^TD^ relative to wild type, yet do not make structured contacts with E2F^TD^: S474R (27-fold); and S474I (27-fold). At 37°C, several additional variants in structured areas outside of the E2F^TD^ binding cleft also weaken Rb-E2F^TD^ binding (**Figure 2F**). These occur within the hydrophobic core of subdomain A (A488V, A490S, A490T, L550I) and the hydrophobic core of subdomain B (M695I, I703F, I703V, M704V), as well as within other regions of the protein fold. Within the hydrophobic core of subdomain B, the variant I703F has a strong temperature dependent effect on weakening E2F1^TD^ binding. For I703F, E2F1^TD^ binding is weakened by approximately 5-fold relative to wild type at 25°C, yet this effect increases to 17-fold at 37°C (**Table 1**, **Figure 2E**, **Figure 2F**). Together these results reveal the large extent of E2F^TD^ binding loss caused by cancer-associated missense variants. This analysis further reveals that E2F^TD^ binding loss is caused by variants located throughout the structured region of Rb^P^ and not limited to those located at the E2F^TD^ binding cleft.

To measure the Rb^P^-E7^LxCxE^ protein-protein interaction, the equilibrium dissociation constant for wild type Rb^P^ binding to TMR-E7^LxCxE^ was measured to be K_D_ = 5 ± 1nM (**Table 1**, **Figure 3A**). This value is similar to the reported K_D_ between Rb^P^ and FITC-E7^LxCxE^ (**Chemes 2010**). When this experiment is conducted with 75 missense variants of Rb^P^, 15% of the variants tested (n=11) have a K_D (app)_ approximately 3-fold, or greater, than the K_D_ for wild type (**Table 1**, **Figure 3B, Supplemental Figure 1**). At 37°C, wild type Rb^P^ binds to E7^LxCxE^ with K_D (app)_ = 12 ± 1nM (**Table 1**, **Figure 3C**). At physiological temperature, 21% of the variants tested (n=16) have a 3-fold, or more, higher K_D (app)_ than the K_D (app)_ for wild type (**Table 1**, **Figure 3D, Supplemental Figure 1**). When the mutation sites are mapped onto the structure of Rb^P^-E7^LxCxE^, all of the mutation sites negatively affecting E7^LxCxE^ binding at 25°C sit within subdomain B, close to the LxCxE binding site (**Figure 3E**). At 37°C, additional sites within subdomain B have greater than 3-fold effects, as do sites outside of subdomain B: E492Q; E554K; and A525G (**Figure 3F**). The biggest reduction in E7^LxCxE^ binding occurs for S751Y, which reduces the Rb^P^-E7^LxCxE^ binding interaction relative to wild type by approximately 26-fold at 25°C and 35-fold at 37°C (**Figure 3E**, **Figure 3F**). The strongest temperature dependent effects reduce Rb^P^-E7^LxCxE^ binding relative to wild type by up to several fold more at physiological temperature vs. standard temperature. This is strongest for I703F, which reduces Rb^P^-E7^LxCxE^ binding approximately 3-fold at 25°C and approximately 23-fold at 37°C (**Table 1**, **Figure 3E**, **Figure 3F**). Because the molecular determinants of Rb^P^-LxCxE-binding interactions are similar across proteins, the mutations identified here as those that weaken Rb^P^-E7^LxCxE^ are likely to also weaken other LxCxE interactions mediating repressive complexes (**Singh 2005**, **Putta 2022**).

**Figure 3.**
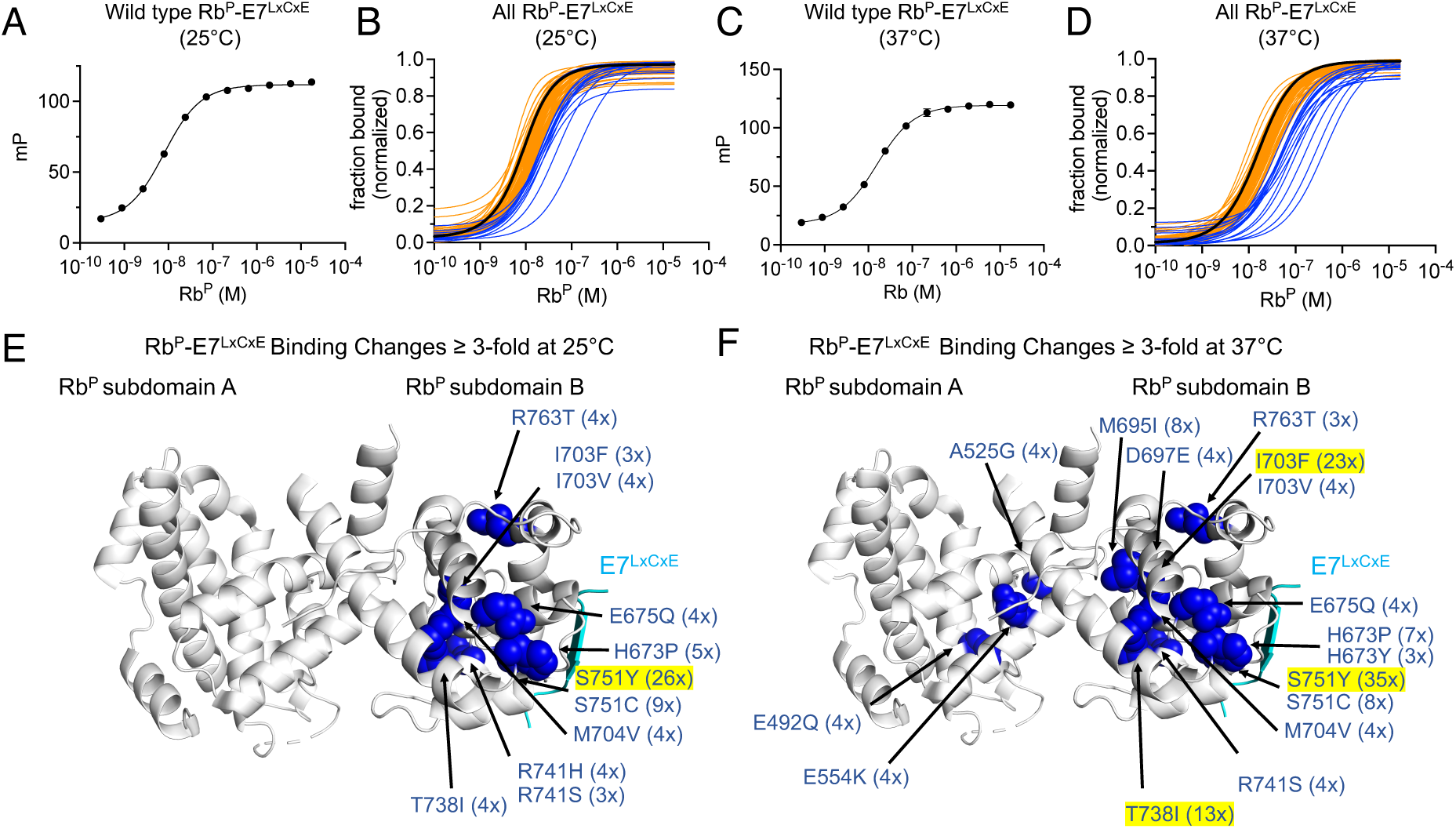
Fluorescence polarization binding experiments for Rb^P^-E7^LxCxE^ at 25°C and 37°C. **A**) The saturating binding curve used to measure binding between TMR-E7^LxCxE^ and wild type Rb^P^ at 25°C. Error bars are standard deviations of four replicates. **B**) Normalized binding curves measured at 25°C depict binding between E7^LxCxE^ and: wild type Rb^P^ (black, n=1); Rb^P^ variants with less than 3-fold increases to K_D (app)_ relative to wild type (orange, n=64); and Rb^P^ variants with 3-fold or more increases to K_D (app)_ relative to wild type (blue, n=11). **C**) The saturating binding curve for TMR-E7^LxCxE^ binding to wild type Rb^P^ at 37°C. Error bars are standard deviations of four replicates. **D**) For binding at 37°C, normalized binding curves are shown for E7^LxCxE^ binding to: wild type Rb (black, n=1); Rb^P^ variants with less than 3-fold increases in K_D (app)_ relative to wild type (orange, n=59), and Rb^P^ variants with 3-fold or more increases in K_D (app)_ relative to wild type (blue, n=16). The structural positions of the amino acid missense variations that show significant increases to K_D (app)_ are mapped onto the known structure of Rb^P^-E7^LxCxE^ for binding experiments conducted at 25°C (**E**) and 37°C (**F**) (blue, space-filling) (PDB: 1GUX) (**Lee 1998**). Fold increases in K_D (app)_ relative to wild type are shown in parentheses next to each mutation name, and those with greater than 10-fold effects are highlighted.

### Protein Instability in Rb^P^ is Caused by Cancer-associated Missense Variants

Wild type Rb and missense variants were evaluated for stability changes by measuring the melting temperature (T_m_), defined as the temperature at which 50% of a protein sample is unfolded (**Huynh 2015**). When wild type Rb^P^ undergoes irreversible melting via Differential Scanning Fluorimetry (DSF), a T_m_ value of 46.9 ± 0.5 °C is calculated from a Boltzmann fit of the melting curve (**Table 1**). This value is similar to reported values from related experiments assessing the stability of Rb pocket domain constructs using circular dichroism: T_m (app)_ = 49°C (**Chemes 2013**) and T_m_ = 46°C (**Huber 1992**). Examination of Rb^P^ missense variants by DSF reveals that approximately 30% (n=30) of all tested variants in both the A and B subdomains have larger than -2°C reductions in T_m_, as measured by the difference between variant T_m_ and wild type T_m_ values. None of the Rb^PL^ variants show greater than -2°C reductions in T_m_ (**Table 1**, **Figure 4A**). This is reasonable because the unstructured pocket loop does not contribute meaningfully to the stability of the folded pocket domain. Within the cyclin fold of subdomain A, the greatest destabilizing effects (ΔT_m_), are caused by E554K (-7.0 ± 0.5 °C), E533K (-6.4 ± 0.5 °C), E492Q (-5.4 ± 0.8 °C °C), A490V (-5.1 ± 0.6 °C), K462E (-5.0 ± 0.5 °C), E539D (-4.9 ± 0.5 °C), A490T (-4.7 ± 1.0 °C) and A488V (-4.1 ± 0.6 °C) (**Table 1**, **Figure 4A**). Within subdomain B, the greatest destabilizing effects are caused by I703F (-6.6°C ± 0.6 °C), S751Y (-5.4 ± 0.5 °C), T738I (-5.3 ± 0.5 °C), M704V (-5.1 ± 0.9 °C) and D697E (-4.1 ± 0.5 °C) (**Table 1**, **Figure 4A**). Examples of raw fluorescence traces used to calculate and evaluate the T_m_ values for wild type Rb^P^, E533K, E554K, I703F, M704V and S751Y are shown (**Figure 4B**). Missense variants that pass a threshold of ΔT_m_ ≤ -3 °C relative to wild type were mapped on to the structure of Rb^P^ (**Figure 4C**). Several of the destabilizing variants occur at amino acid sites completely buried within the hydrophobic cores of subdomain A (A488, A490, I550) and subdomain B (M695, I703, M704). From the structure map, it is also apparent that all of the mutations which cause destabilizing effects of ΔT_m_ ≤ -3 °C also weaken Rb^P^ binding to E2F1^TD^ by 3-fold or more (**Figure 2F and Figure 4C**).

**Figure 4.**
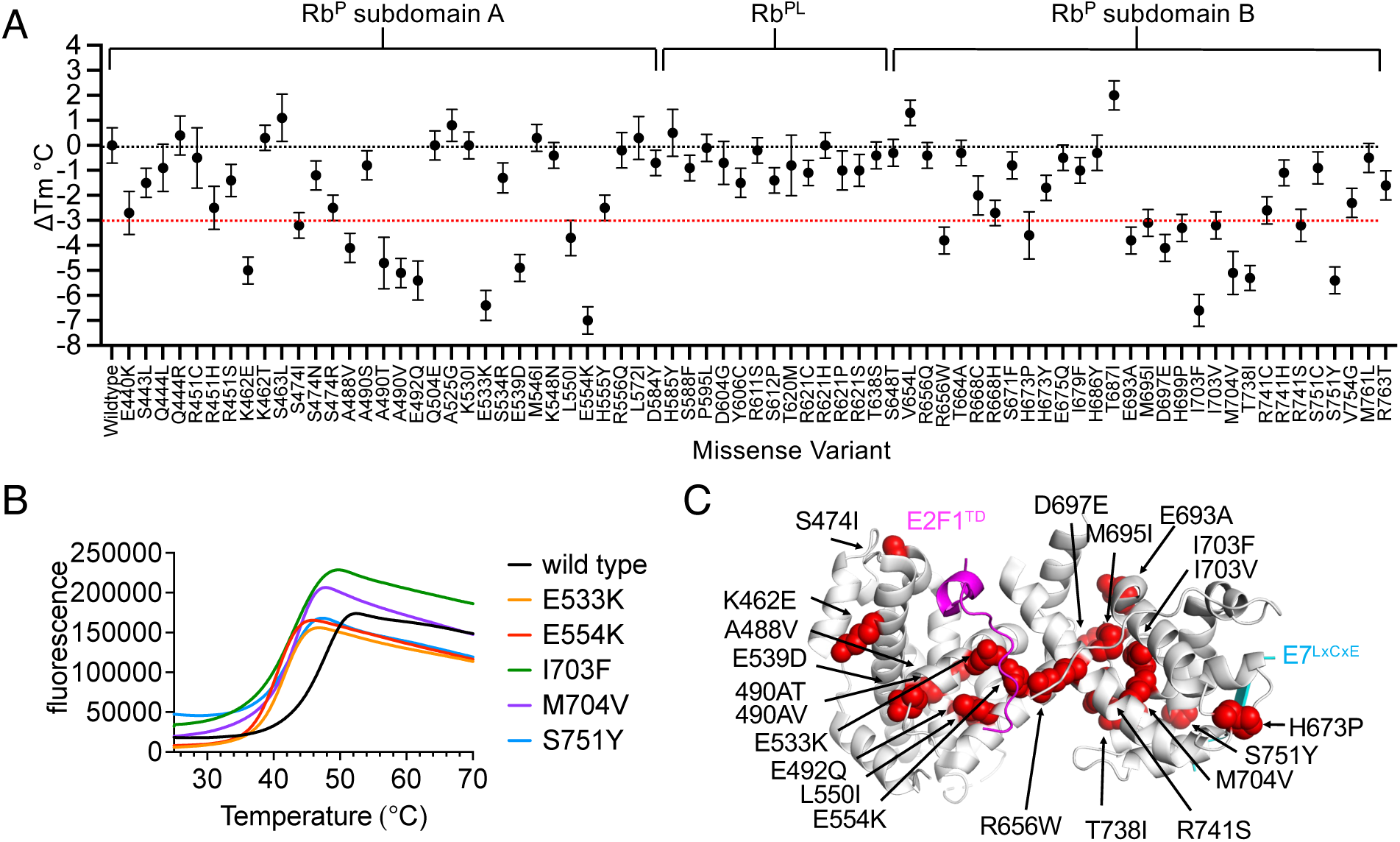
Changes to the thermostability of Rb^P^ caused by missense variants. **A**) Differences between the T_m_ values of Rb^P^ variants relative to wild type Rb^P^ (ΔT_m_) reveal the relative size of destabilizing effects caused by each variant. Error bars are propagated from standard deviations of averages calculated from 9 replicates for wild type and each missense variant. **B)** Examples of raw fluorescence data for DSF runs of wild type Rb^P^ and four example variants which show some of the greatest destabilizing effects. **C)** Missense variants that cause a negative shift in T_m_ greater than a 3°C threshold mapped onto the structure of Rb^P^.

### Relationship Between T_m_ and Temperature Sensitive Binding for Rb^P^ Variants

To explore the possibility that missense variants which cause greater destabilizing effects also disrupt binding to E2F1^TD^ more at higher temperatures, a plot was made of ΔT_m_ values (x-axis) *vs*. the ratio of measured K_D (app)_ values for Rb^P^-E2F1^TD^ at 37°C and 25°C (y-axis) (**Figure 5A**). A linear fit of this scatter plot reveals a positive association between the -ΔT_m_ change and the quotient of K_D (app)_ 37°C / K_D (app)_ 25°C. A similar relationship is also seen for Rb^P^-E7^LxCxE^ (**Figure 5B**). From this analysis, the variant I703F stands out as being the most temperature sensitive for both E2F1^TD^ and E7^LxCxE^ binding; however, variants from other regions of the protein, E554K and S751Y, also follow the trend. In general, these relationships may be useful to predict variants that are temperature sensitive based only on their T_m_ values, or alternatively, multitemperature binding data may be used identify variants that are likely destabilizing. This approach may be particularly useful for identifying cryptic disease-associated variants that are less thermostable yet do not significantly weaken binding interactions at 25°C; since these may be more likely to show effects at 37°C.

**Figure 5.**
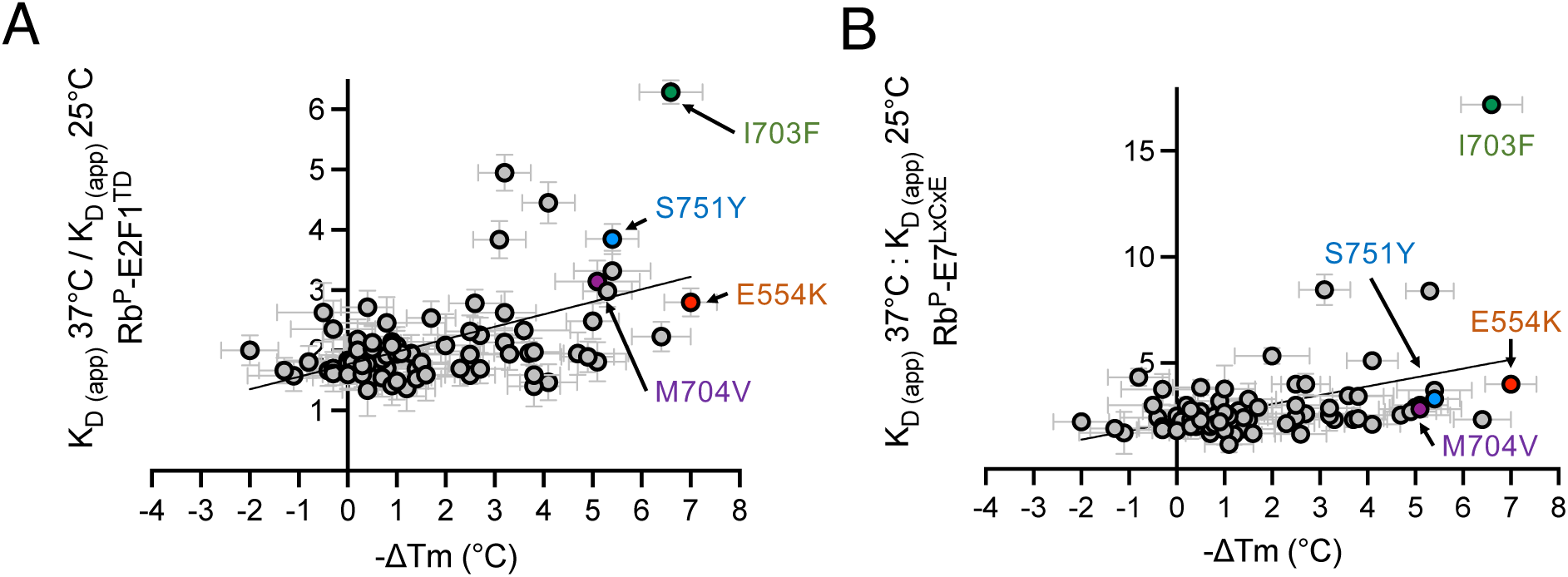
Relationship between ΔT_m_ and temperature sensitive binding. Scatter plot of the ratio of K_D (app)_ values (37°C: 25°C) for E2F1^TD^ (**A**) and E7^LxCxE^ (**B**), as a function of -ΔT_m_. Error bars are propagated from FP standard deviations and DSF standard deviations between 4 and 9 replicates, respectively. The data were fit to a linear regression: y=0.04081x+0.1125 with r^2^=0.26 for (**A**) and y=0.1073x+0.2958 with r^2^=0.17 for (**B**).

One explanation for the observed relationship between the measured ΔT_m_ values and the temperature sensitive binding effects is that less stable variants become irreversibly unfolded at higher temperatures, creating conditions that appear as weaker binding. To explore this, four variants were selected from different regions of Rb^P^ to examine the reversibility of temperature dependent effects of E2F1^TD^ and E7^LxCxE^ binding. For each variant, the K_D (app)_ is measured initially at 25°C, then a after temperature ramping and incubating at 37°C, the K_D (app)_ is measured again. A third measurement of K_D (app)_ is taken of a sample that is ramped to 37°C, incubated at 37°C, then cooled down again to 25°C. The saturating binding curves produced under these 3 conditions reveal that in each case, the pre and post warming measurements at 25°C are each very close to one another, while the measurement at 37°C is significantly different (**Figure 6A-J**). The K_D (app)_ values for Rb^P^ binding to E2F1^TD^ at 25°C, for the third measurement, measured after warming to 37°C, then cooling to 25°C are: 17 ± 5 nM for wild type; 1737 ± 224 nM for E554K; 94 ± 13 nM for I703F; 46 ± 9 nM for M704V; and 28 ± 8 nM for S751Y. The K_D (app)_ values for Rb^P^ binding to E7^LxCxE^ at 25°C, measured after warming and cooling are: 7 ± 1 nM for wild type; 24 ± 2 nM for E554K; 53 ± 6 nM for I703F; 14 ± 1 nM for M704V; and 12 ± 1 nM for S751Y. Reported errors in the K_D (app)_ measurements are curve-fitting errors. The other values for K_D (app)_, measured at 25°C and 37°C, shown in **Figure 6A-J** are given in **Table 1**. Together these results show that weaker binding at 37°C is completely or nearly completely reversible for wild type, E554K, M704V and S751Y, as tested under these conditions. The exception is I703F, which upon cooling back down to 25°C from 37°C, binds significantly weaker to E7^LxCxE^ (K_D (app)_ 53 ± 6 nM) than it does when initially measured at 25°C (K_D (app)_ 16 ± 2 nM). The data suggest that reversible fold destabilization or alternative fold conformations may underly the temperature sensitivity of some missense variants, while others irreversibly unfold at physiological temperature. Future studies of binding enthalpies will be needed to provide more complete descriptions of thermodynamic relationships underlying the effects of Rb^P^ variants.

**Figure 6.**
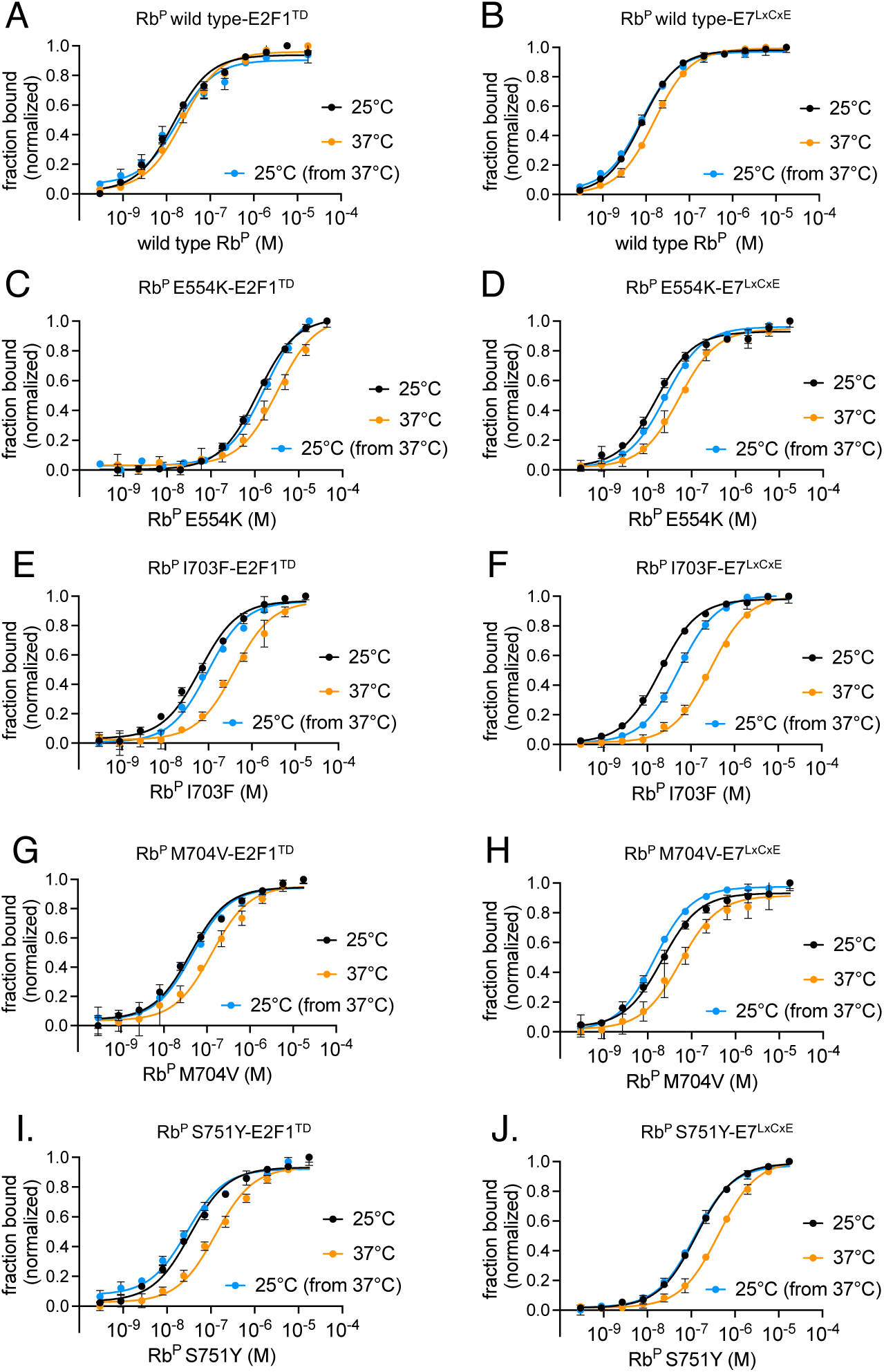
Reversibility of temperature sensitive binding effects for wild type, E554K, I703F, M704V and S751Y. Fluorescence polarization experiments were performed to assess the reversibility of temperature dependent binding changes to E2F1^TD^ and E7^LxCxE^ for Rb^P^ wild type (**A, B**), E554K (**C, D**), I703F (**E, F**), M704V (**G, H**), S751Y (**I, J**). The experiments measured binding initially at 25°C (black), then at 37°C (orange), and also after cooling down to 25°C from 37°C (blue). Error bars are standard deviations of four replicates.

### X-ray Crystal Structures Reveal Causes of Cancer Mutation Effects

For characterization by protein X-ray crystallography, four variants were studied: E533K, E554K, M704V and S751Y. In order to crystalize these variants, each missense mutation was independently introduced into a modified Rb^P^ construct, previously used to crystalize the Rb^PL^-Rb^P^ interaction (**Burke 2012**). X-ray diffraction data collected from the crystals were used to solve the structures of the four variants by molecular replacement (**Table 2**). These specific variants were selected because they cause some of the largest measured decreases in thermostability (ΔT_m_ ≤ -5 °C), strongly disrupt E2F1^TD^ or E7^LxCxE^ binding, are temperature sensitive, and map to key regions of the pocket domain (**Figure 7A**).

**Figure 7.**
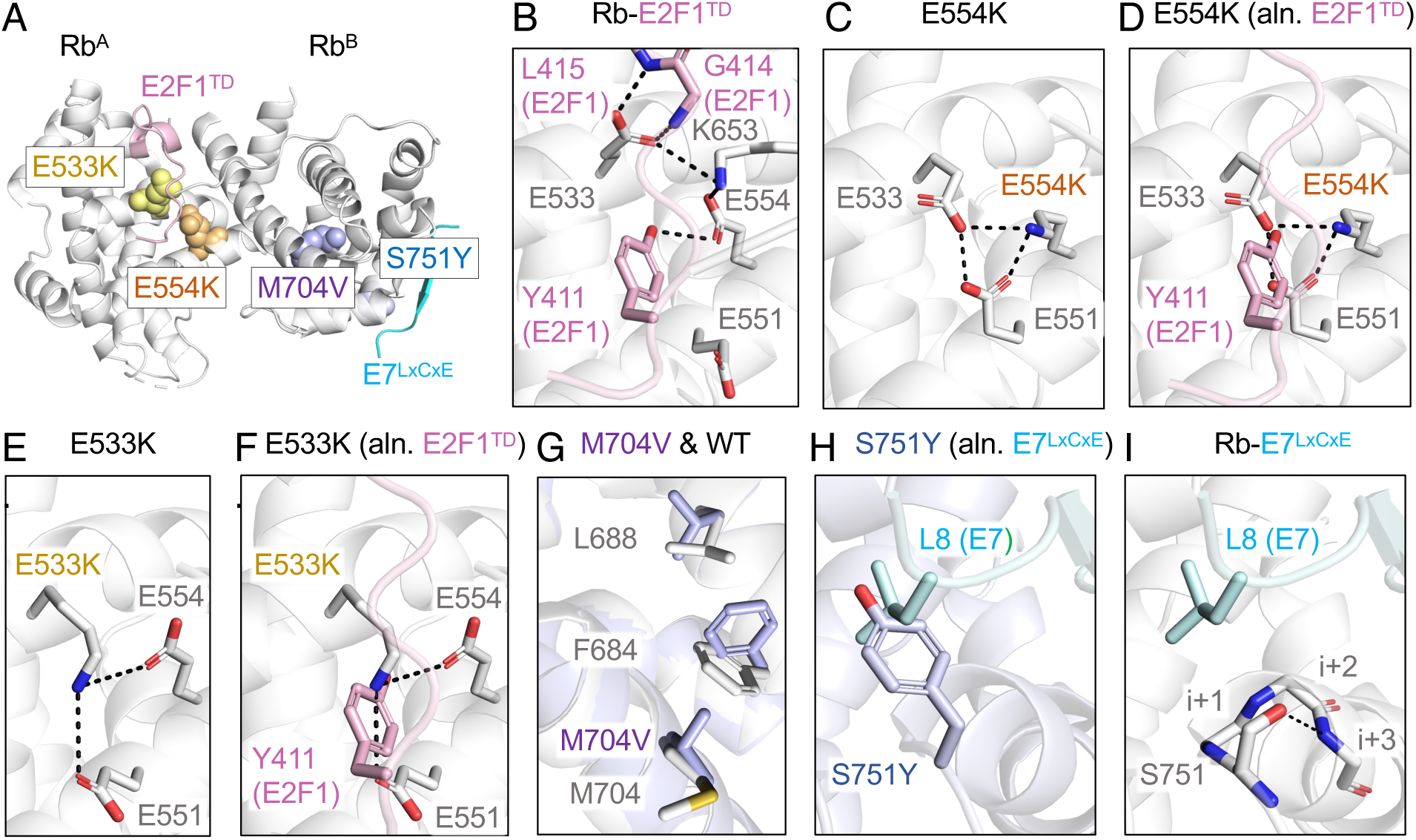
Protein X-ray crystal structures of missense variants and comparisons to wild type Rb^P^ structures. **A)** The locations of crystallized variants in the pocket domain relative to E2F1^TD^ and E7^LxCxE^ binding sites. **B)** Wild type Rb^P^-E2F1^TD^ biding interactions are mediated by E533 and E554, from a previously published structure (PDB: 1O9K) (**Lee 2002**). **C)** The structure of E554K shows the formation of a salt bridge network involving E554K, E533 and E551. **D)** A structural alignment of the wild type Rb^P^-E2F1^TD^ with the structure of E554K reveals that the E554K-mediated salt bridge network blocks an important tyrosine (Y411) binding site of E2F^TD^. **E)** The structure of E533K shows the formation of a salt bridge network involving E533K, E554 and E551, similar to the E554K structure. **F)** A structural alignment of the wild type Rb^P^-E2F1^TD^ with the structure of E533K reveals that the salt bridge network created by the missense variant blocks the tyrosine (Y411) binding site of E2F^TD^, similar to the effect of E554K. **G)** The structure of M704V reveals how the mutation induces a change in the packing of the hydrophobic core of subdomain B. **H)** The structure of S751Y aligned to the structure of Rb^P^-E7^LxCxE^ shows how the tyrosine variant occupies the binding pocket for L8 in of E7^LxCxE^ peptide (PDB: 1GUX) (**Lee 1998**). **G)** The previously published structure of wild type Rb^P^-E7^LxCxE^ reveals S751 forms an H-bond to stabilize an ⍺-helix, the loss of which may contribute to the reduced stability of S751Y (PDB: 1GUX) (**Lee 1998**).

**Table 2.** X-ray crystallography data collection and refinement statistics. Parentheses are for the highest resolution shell.

|  | Rb E533K | Rb E554K | Rb M704V | Rb S751Y |
| --- | --- | --- | --- | --- |
| **Data collection** |  |  |  |  |
| Space group | R 3 :H | R 3 :H | R 3 :H | R 3 :H |
| Cell dimensions |  |  |  |  |
| *a*, *b*, *c* (Å) | 250.14, 250.14, 35.31 | 250.36, 250.36, 35.31 | 250.84, 250.84, 35.39 | 255.04, 255.04, 35.36 |
| $\alpha$, $\beta$, $\gamma$ (°) | 90, 90, 120 | 90, 90, 120 | 90, 90, 120 | 90, 90, 120 |
| Resolution (Å) | 47.27 – 2.16 | 47.31 – 2.26 | 47.4 – 2.38 | 48.2 – 2.32 |
| *R*<sub>merge</sub> | 0.191 (2.665) | 0.215 (2.575) | 0.233 (3.180) | 0.174 (2.774) |
| *R*<sub>pim</sub> | 0.092 (1.284) | 0.104 (1.301) | 0.079 (1.130) | 0.085 (1.382) |
| CC ½ | 0.997 (0.401) | 0.997 (0.369) | 0.997 (0.381) | 0.998 (0.302) |
| *I* / σ*I* | 9.9 (1.1) | 9.6 (1.0) | 10.2 (0.9) | 12.4 (0.9) |
| Completeness (%) | 100.0 | 100.0 | 100.0 | 100.0 |
| Multiplicity | 10.4 | 10.4 | 10.5 | 10.5 |
| **Refinement** |  |  |  |  |
| Resolution (Å) | 41.69 – 2.16<br>(2.237 – 2.16) | 41.73 – 2.26<br>(2.314 – 2.26) | 34.78 – 2.38<br>(2.465 – 2.38) | 48.2 – 2.32<br>(2.403 – 2.32) |
| No. unique reflections | 44174 (4393) | 38642 (3919) | 33256 (3275) | 37130 (3729) |
| *R*<sub>work</sub> / *R*<sub>free</sub> | 0.2260 (0.3148) /<br>0.2623 (0.3373) | 0.2273 (0.3286) /<br>0.2653 (0.3725) | 0.2203 (0.3313) /<br>0.2772 (0.3737) | 0.2126 (0.3641) /<br>0.2629 (0.4060) |
| No. atoms | 5753 | 5791 | 5694 | 5793 |
| Protein | 5679 | 5653 | 5685 | 5715 |
| Water | 74 | 138 | 9 | 78 |
| B-factors (Å<sup>2</sup>) |  |  |  |  |
| Average | 66.51 | 58.99 | 71.70 | 76.89 |
| Protein | 66.68 | 59.24 | 71.71 | 77.24 |
| Water | 53.25 | 48.61 | 62.82 | 50.88 |
| R.m.s. deviations |  |  |  |  |
| Bond lengths (Å) | 0.015 | 0.016 | 0.015 | 0.015 |
| Bond angles (°) | 1.50 | 1.41 | 1.51 | 1.50 |
| Ramachandran Plot |  |  |  |  |
| Favored (%) | 98.51 | 96.71 | 96.73 | 97.33 |
| Allowed (%) | 1.49 | 3.14 | 3.12 | 2.67 |
| Outliers (%) | 0.00 | 0.15 | 0.15 | 0.00 |
| PDB codes | 9DHU | 9DHF | 9DGK | 9DHC |

For E554K and E533K, we sought to understand the structural basis for large reductions in Rb^P^-E2F1^TD^ binding as well as substantial decreases in fold stability. The significance of E554 in Rb-E2F^TD^ binding was pointed out initially by the authors of the two Rb^P^-E2F^TD^ structures; specifically, it forms an H-bond the phenolic hydroxyl of conserved tyrosine in E2F1^TD^ and E2F2^TD^ (**Lee 2002, Xiao 2003**) (**Figure 7B**). Additionally, from the structure of Rb^P^-E2F1^TD^, it can be seen that the sidechain of E553 forms two H-bonds with the backbone amides of L415 and G414 of E2F1^TD^ (**Figure 7B**). These structured interactions show the roles of glutamate sidechains of E553 and E554 in mediating binding interactions between wild type Rb^P^ and E2F^TD^. The structure of the cancer-associated variant, E554K, reveals how a lysine substitution disrupts the potential for these interactions by forming a shared salt bridge between E553 and E551 (**Figure 7C**). This high-energy intramolecular bond does two notable things that likely disrupt E2F^TD^ binding: 1) it stabilizes a rotamer of E533 in an orientation is not amenable to E2F^TD^ backbone binding, and 2) it places E533 and E551 in a triad with E554K that sterically occludes the binding pocket occupied by Y411 (**Figure 7D**). Remarkably, the crystal structure of E533K shows something similar: the mutated lysine sidechain creates a salt bridge triad with E554 and E551 in a manner that occludes the Y411(E2F1^TD^) binding pocket (**Figure 7E**, **Figure 7F**). Together the variant structures of E533K and E554K reveal a structural basis for the disruption of E2F^TD^ binding. However, from these structures alone it is not clear how the mutations induce fold destabilization. In the structures of Rb^P^-E2F2^TD^ Rb^P^-E2F1^TD^, as well as in the unliganded “apo” Rb^P^ structure, both E554 and E533 form A/B subdomain-spanning salt bridges to K653 (**Lee 2002, Xiao 2003, Balog 2011**) (**Figure 7B, Supplemental Figure 2**). It is possible that the loss of the salt bridge interactions contributes to the large destabilization effects observed for these variants (**Figure 4A-C**).

Missense variants within the hydrophobic core of subdomain B, such as I703F, I703V, M695I and M704V, show reduced stability and temperature sensitive binding to E2F1^TD^ and E7^LxCxE^ (**Figure 5A**, **Figure 5B**). To understand how these mutations may cause these effects, we crystallized M704V. The crystal structure of M704V reveals changes to the packing of the hydrophobic core of subdomain B (**Figure 7G**). Specifically, M704V introduces new steric bulk in the form of the valine gamma carbon. This forces alternative rotamers of F684 and L688 through changes to a van der Waals contact network. These specific changes to sidechain orientations of F684 and L688 are not observed in any other structure of Rb^P^. This structure therefore provides some insight into how M704V destabilizes Rb^P^ and the basis for temperature sensitive binding to E2F1^TD^.

For a structural example of E7^LxCxE^ inhibition, S751Y was crystallized. This variant weakens both E2F1^TD^ and E7^LxCxE^ binding relative to wild type, but causes the greatest reduction to E7^LxCxE^ binding of all missense variants tested here. The structure of S751Y reveals that the tyrosine sidechain of the mutant sits within a hydrophobic binding pocket used for LxCxE binding (**Figure 7H**). Specifically, the tyrosine sits in a position to compete for the binding of a conserved I or L, which constitutes a significant part of the LxCxEx(I/L) binding epitope (**Singh 2005**, **Putta 2022**). Even though the variant tyrosine creates a snug fit into the pocket used for I/L binding, this variant is not stabilizing; instead, it causes a large destabilizing effect (ΔT_m_ = -5.4 ± 0.5 °C). In the structure of wild type Rb-E7^LxCxE^, the S751 sidechain helps to stabilize the N-terminus of an alpha helix through an ST motif-stabilizing H-bond to the amide at i+3 (**Lee, 1998, Wan 1999**) (**Figure 7I**). Loss of this known helix-stabilizing motif may be responsible for the large thermostability decrease seen for S751Y.

In summary, these crystal structures reveal several novel and unexpected insights into the structural changes that help to support the related biochemical findings. Overall, this work provides a comprehensive and detailed understanding of both structural and biochemical outcomes of specific variants. The information provided by this study will be useful for informing predictions of the potential broader consequences of these and of yet unstudied variants.

## DISCUSSION

Cancer genome sequencing has revealed detailed landscapes of somatic missense mutations that can provide insights to various mechanisms of inactivation of tumor suppressor proteins (**Cancer Genome Atlas Research Network 2013**). Here we select many of the recurrent mutations to the pocket domain of Rb and use biochemical methods to quantify changes to key binding interactions and thermostabilities, as well as structural changes caused by these variants through protein X-ray crystallography. The most significant finding of this data set is the identification of cancer associated mutations causing disruptions in binding to E2F^TD^; a critical interaction with strong connections to cancer cell cycles (**Rubin 2020**). Surprisingly, we found that nearly half of the missense variants tested (32/75) showed 3-fold or greater E2F1^TD^ binding inhibition at physiological temperature. Variants with the strongest effects map to the E2F^TD^ binding cleft, but others throughout the A and B subdomains show effects as well. When the same variants were examined for disruptions in binding to E7^LxCxE^, 16 variants show 3-fold, or greater, binding inhibition at physiological temperature; these are located primarily on the B subdomain. There is considerable overlap between the variants that affect E7^LxCxE^ binding with those that also affect E2F1^TD^ binding, and these also tend to be the variants which reduce the thermostability of Rb^P^. There are also mutations that don’t have overlapping effects. Specifically, out of the 75 variants tested only 4 selectively inhibited E7^LxCxE^ and not E2F1^TD^ and these effects were all under 5-fold: A525G; R741C; R741S; and R763T. On the other hand, several variants strongly inhibit E2F1^TD^, by greater than 10-fold, but do not affect E7: S534R; S474I; S474R; K530I; H555Y; and R656Q. Some of these variants have been observed in clinical studies. The variant H555Y was identified in a multiplexed immunofluorescent approach examining subclonal lung adenocarcinoma tumor populations using 5-cell profiles; however, the authors could not justify the clinical significance of finding this mutant (**Zhu 2022**). Two incidents of the variant R656Q were identified from a genetic screen of 150 retinoblastoma patients including one unilateral case and one bilateral case from unrelated patients (**Sagi 2015**).

Protein X-ray crystallography revealed structural bases for destabilization and binding defects for key variants: E533K; E554K; S751Y and M704V. Although, it was not clear from the structure of M704V why E2F1^TD^ and E7^LxCxE^ binding are affected by this variant, since it is distant from binding sites. A similar crystallographic study on p53 revealed a mutation known to have temperature sensitive DNA binding effects displays only subtle changes to sidechain packing within the hydrophobic core of the p53 core-domain (**Wallentine 2013**). While Rb’s M704V mutation alters the hydrophobic core’s energetic ground state by reconfiguring several buried residues, these changes must also have structural consequences that extend beyond the hydrophobic core. Additional experiments will be needed to elucidate the structural mechanisms through which changes to core packing translate to E2F^TD^ and E7^LxCxE^ binding inhibition by Rb^P^ hydrophobic core variants.

Many of the variants that we attempted to study did not express in *e coli*. While this may seem to limit the scope of this study, we consider the variants that were measured in this study significant because they exist in the context of a folded pocket domain, and therefore any major effects measured for these variants are not due to irreversibly unfolded protein, but rather to specific binding changes that presumably leave other Rb protein functions intact. We were not able to use our methods to study the most commonly observed, pathological Rb^P^ missense mutation, R661W. However, R661W has successfully expressed in other host systems where intact binding-dependent functions provide evidence of its folding (**Otterson 1999, Cobrinik 2006**). Of the temperature sensitive variants that we were able to study, I703F displayed the strongest effect, reducing binding 6-fold more at physiological temperature (37°C) compared with room temperature (25°C) for E2F1^TD^. For E7^LxCxE^, 16-fold weaker binding was measured at physiological temperature relative to room temperature. Significantly, this was also the most destabilizing variant studied, as it reduced the T_m_ of the protein 6.6°C relative to wild type. Under the binding conditions at physiological temperatures, the temperature sensitivity for the few variants tested were reversible; however, it’s very likely that under extended incubations at physiological temperature, irreversible unfolding of the pocket domain may be a predominant mechanism for loss of binding activity for many variants. Stability studies on wild type Rb pocket have shown how it is susceptible to irreversible unfolding *in vitro*, such that 50% of the protein is unfolded in 30 minutes at physiological temperature (**Chemes 2013**). On the other hand, E2F1^TD^ and E7^LxCxE^ binding to Rb^P^ both produce stabilizing effects for Rb^P^ (**Huber 1992, Chemes 2013**). Taking this into consideration with our findings, it’s likely that some missense mutations in the cancer cell have dual destabilizing effects on Rb^P^: they destabilize Rb^P^ directly yet also further destabilize Rb^P^ through disruptions to endogenous interactions at the E2F^TD^-binding site and the LxCxE-binding site.

## METHODS

### Cloning, protein expression and purification

Missense variants were cloned into a human Rb^P^ gene construct (corresponding to amino acids 380-788) (UniProt P06400) in a pGEX-4T expression vector (GenBank M21676, M97937). Primers were designed using an optimized QuickChange protocol for site-directed mutagenesis (**Supplemental Table 1**) (**Liu 2008**) and ordered through a commercial vendor (IDT). Plasmids were sequenced to confirm the presence of the mutations (Plasmidsaurus, Genewiz). Plasmids were transformed into BL21 DE3-pRIL modified *e coli* cells, grown in Terrific broth, and induced overnight with 1 mM IPTG at 25°C. Cells were lysed using sonication and cell debris was pelleted through centrifugation. The overexpressed GST fusions in the supernatant were purified by glutathione affinity chromatography in a buffer containing 200 mM NaCl, 1 mM Dithiothreitol (DTT), and 25 mM Tris–HCl (pH 8.0) and eluted using 10 mM glutathione. The fusion proteins were cleaved from the GST affinity tag using 1% TEV protease at 4°C overnight. Proteins were then further purified by cation exchange chromatography on a heparin column in tandem with a GST trap, in order to collect residual uncut GST-Rb or GST tag. Protein purity was characterized by SDS-PAGE and UV 260/280.

### Fluorescence Polarization

FP experiments were set up using Matrix Equalizer electronic multichannel pipettes in 20 μl reaction volumes in black untreated 384-well plates (Corning). All solutions were prepared using a buffer containing 25 mM Tris (pH 8.0), 200 mM NaCl, 1 mM DTT, and 0.1% Tween-20. Synthetic, fluorescently-labeled peptides were ordered from GenScript: TMR-E7^LxCxE^ (DLYCYEQLN), TMR-E2F1^TD^ (GEGIRDLFDCDFGDLTPLD). For the binding experiments, concentrations of 1.5 nM TMR-E2F1^TD^ and 7.5 nM TMR-E7^LxCxE^ were used with unlabeled Rb^P^. For the 25°C experiment, samples were allowed to incubate for two minutes at 25°C to ensure that binding equilibrium had been reached. Fluorescence polarization was measured on a PHERAstar Microplate Reader using Labtech software (BMG). Reported y-axis values of millipolarization (mP) were determined from the software and calculated from the relationship: mP=1,000 × (S−G × P)/(S+G × P), such that S is the fluorescence intensity parallel to the excitation plane, P is the perpendicular fluorescence intensity and G is the correction factor. For the 37°C experiment: following measurement at 25°C, the temperature of the PHERAstar instrument (with sample plate) was ramped up over a duration of 7 minutes to 37°C, followed by a 2-minute incubation at 37°C before FP data was collected. For the 37°C back to 25°C experiment, sample plates were incubated at 25°C, ramped up to 37°C in an Heratherm plate incubator (ThermoFisher) over 7 minutes with careful temperature monitoring by thermometer, incubated for 2 minutes at 37°C, cooled on ice for 1 minute, then incubated at 25°C for three minutes prior to taking FP measurements. All experiments were conducted in quadruplicate. A quadratic equation was used to fit the binding data and calculate the K_D_ or apparent K_D_ in Prism 9 (GraphPad) (**Jarmoskaite 2020**). Reported errors in the K_D_ or apparent K_D_ from FP measurements are curve-fitting errors (confidence interval 95%) and error bars are standard deviations of four replicates. Data was collected and processed using BMG Labtech and MARS software (BMG).

### Differential Scanning Fluorimetry

DSF assays were performed using SYPRO orange dye (ThermoFisher Scientific) in a 1:625 dilution in a buffer containing 200 mM NaCl, 25 mM Tris (pH 8.0), 2 mM DTT. Reactions were run in 20 μl reaction volumes with final Rb^P^ concentrations of 3 μM, 6 μM and 9 μM. Replicates of three experiments were conducted for each condition. Experiments with different dye to protein ratios were averaged together in order to average out concentration dependent effects (**Huynh 2015**). All experiments were performed on a QuantStudio 3 Real-Time PCR (ThermoFisher Scientific) and the raw fluorescence data was fit using a Boltzmann function to calculate the melting temperature (T_m_) on Protein ThermalShift software (ThermoFisher Scientific). Reported T_m_ values are the average values for the T_m_ across all replicates across conditions and the reported error is the calculated standard deviation. Change in melting temperature values (ΔT_m_) were calculated by subtracting variant T_m_ from wild type T_m_ values.

### Protein crystallization, data collection, and processing

Missense variants of Rb^P^ characterized by protein X-ray crystallography were first cloned into a Rb^P^ construct (Rb^380–787D616–642/S608E/S612A/S780A^) previously used to crystallize the Rb^PL^-Rb^P^ interaction (**Burke 2012**). The E533K, E554K, M704V and S751Y missense mutations were introduced to this specific construct because it crystalizes reproducibly. Site-directed mutagenesis was accomplished using a modified QuickChange protocol (**Liu 2008**). Proteins were expressed from *e. coli* and purified as described previously, however for this application they were also further purified by size exclusion chromatography on a Sephadex 200 column, and with a buffer containing 200 mM NaCl, 25 mM Tris–HCl (pH 8.0) and 2 mM DTT. Eluate fractions were concentrated to approximately 7 mg/ml and crystals were grown using hanging drop diffusion at 4°C for 1 week in a 1:1 ratio with a condition containing 18% Peg 8K, 0.1 M sodium citrate, 0.1M succinate pH 5.5, and 1M lithium chloride. Crystals were cryogenically stabilized in a solution containing 20% glycerol and flash frozen in liquid nitrogen. X-ray diffraction data was collected at ALS BL5.0.1. Data were indexed and integrated with XDS (**Kabsch 2010**) in the space group R3:H and merged and scaled with AIMLESS (**Evans 2011**). Molecular replacement solutions were found using PHASER (**McCoy 2007**) with the search model PDB ID: 4ELL (**Burke 2012**). Atomic models were built and modified using COOT (**Emsley 2010**), and refined using default parameters in PHENIX (**Liebschner 2019**). Structure coordinates and maps were deposited to the Protein Data Bank with the following codes: Rb E533K (9DHU); Rb E554K (9DHF); Rb M704V (9DGK); Rb S751Y (9DHC).

## Acknowledgments

We thank the staff at ALS BL 5.0.1 for assistance and support in collecting data.

## Author contributions

Conceptualization: JRB

Methodology: AC, ERW-S, SMR, JB

Investigation: AC, AR-R, CCM, ERW-S, HNM, VIV-M, MAR, MML, IB, SBB, EV, ST, JRB

Visualization: AC, JRB

Supervision: JRB Writing—original draft: JRB

Writing—review & editing: AC, AR-R, CCM, ERW-S, HNM, VIV-M, MAR, MML, IB, SBB, EV, ST, JRB

## Data and materials availability

All data are available in the main text or the supplementary materials. Protein X ray structures have been deposited and are available from the Protein Data Bank under the following accession numbers: Rb E533K (PDB: 9DHU) Rb E554K (PDB: 9DHF) Rb M704V (PDB: 9DGK) Rb S751Y (PDB: 9DHC)

## REFERENCES

1. Altschuler SE, Lewis KA, Wuttke DS. Practical strategies for the evaluation of high-affinity protein/nucleic acid interactions. J Nucleic Acids Investig. 2013 Jan 1;4(1):19–28. doi: 10.4081/jnai.2013.e3. PMID: 25197549; PMCID: PMC4155600.

2. Balog ER, Burke JR, Hura GL, Rubin SM. Crystal structure of the unliganded retinoblastoma protein pocket domain. Proteins. 2011 Jun;79(6):2010–4. doi: 10.1002/prot.23007. Epub 2011 Apr 12. PMID: 21491492; PMCID: PMC3092862.

3. Bourgo RJ, Thangavel C, Ertel A, Bergseid J, McClendon AK, Wilkens L, Witkiewicz AK, Wang JY, Knudsen ES. RB restricts DNA damage-initiated tumorigenesis through an LXCXE-dependent mechanism of transcriptional control. Mol Cell. 2011 Aug 19;43(4):663–72. doi: 10.1016/j.molcel.2011.06.029. PMID: 21855804; PMCID: PMC4271833.

4. Burke JR, Deshong AJ, Pelton JG, Rubin SM. Phosphorylation-induced conformational changes in the retinoblastoma protein inhibit E2F transactivation domain binding. J Biol Chem. 2010 May 21;285(21):16286–93. doi: 10.1074/jbc.M110.108167. Epub 2010 Mar 11. PMID: 20223825; PMCID: PMC2871496.

5. Burke JR, Hura GL, Rubin SM. Structures of inactive retinoblastoma protein reveal multiple mechanisms for cell cycle control. Genes Dev. 2012 Jun 1;26(11):1156–66. doi: 10.1101/gad.189837.112. Epub 2012 May 8. PMID: 22569856; PMCID: PMC3371405.

6. Burke JR, Liban TJ, Restrepo T, Lee HW, Rubin SM. Multiple mechanisms for E2F binding inhibition by phosphorylation of the retinoblastoma protein C-terminal domain. J Mol Biol. 2014 Jan 9;426(1):245–55. doi: 10.1016/j.jmb.2013.09.031. Epub 2013 Oct 5. PMID: 24103329; PMCID: PMC3872205.

7. Burkhart DL, Sage J. Cellular mechanisms of tumour suppression by the retinoblastoma gene. Nat Rev Cancer. 2008 Sep;8(9):671–82. doi: 10.1038/nrc2399. PMID: 18650841; PMCID: PMC6996492.

8. Cancer Genome Atlas Research Network; Weinstein JN, Collisson EA, Mills GB, Shaw KR, Ozenberger BA, Ellrott K, Shmulevich I, Sander C, Stuart JM. The Cancer Genome Atlas Pan-Cancer analysis project. Nat Genet. 2013 Oct;45(10):1113-20. doi: 10.1038/ng.2764. PMID: 24071849; PMCID: PMC3919969.

9. Cerami E, Gao J, Dogrusoz U, Gross BE, Sumer SO, Aksoy BA, Jacobsen A, Byrne CJ, Heuer ML, Larsson E, Antipin Y, Reva B, Goldberg AP, Sander C, Schultz N. The cBio cancer genomics portal: an open platform for exploring multidimensional cancer genomics data. Cancer Discov. 2012 May;2(5):401–4. doi: 10.1158/2159-8290.CD-12-0095. Erratum in: Cancer Discov. 2012 Oct;2(10):960. PMID: 22588877; PMCID: PMC3956037.

10. Chemes LB, Sánchez IE, Smal C, de Prat-Gay G. Targeting mechanism of the retinoblastoma tumor suppressor by a prototypical viral oncoprotein. Structural modularity, intrinsic disorder and phosphorylation of human papillomavirus E7. FEBS J. 2010 Feb;277(4):973-88. doi: 10.1111/j.1742-4658.2009.07540.x. Epub 2010 Jan 20. PMID: 20088881.

11. Chemes LB, Noval MG, Sánchez IE, de Prat-Gay G. Folding of a cyclin box: linking multitarget binding to marginal stability, oligomerization, and aggregation of the retinoblastoma tumor suppressor AB pocket domain. J Biol Chem. 2013 Jun 28;288(26):18923–38. doi: 10.1074/jbc.M113.467316. Epub 2013 Apr 30. PMID: 23632018; PMCID: PMC3696668.

12. Cobrinik D, Francis RO, Abramson DH, Lee TC. Rb induces a proliferative arrest and curtails Brn-2 expression in retinoblastoma cells. Mol Cancer. 2006 Dec 12;5:72. doi: 10.1186/1476-4598-5-72. PMID: 17163992; PMCID: PMC1764425.

13. Cobrinik D. Retinoblastoma Origins and Destinations. N Engl J Med. 2024 Apr 18;390(15):1408-1419. doi: 10.1056/NEJMra1803083. PMID: 38631004.

14. Dick FA, Rubin SM. Molecular mechanisms underlying RB protein function. Nat Rev Mol Cell Biol. 2013 May;14(5):297–306. doi: 10.1038/nrm3567. Epub 2013 Apr 18. PMID: 23594950; PMCID: PMC4754300.

15. Dick FA, Goodrich DW, Sage J, Dyson NJ. Non-canonical functions of the RB protein in cancer. Nat Rev Cancer. 2018 Jul;18(7):442–451. doi: 10.1038/s41568-018-0008-5. PMID: 29692417; PMCID: PMC6693677.

16. Dyson NJ. RB1: a prototype tumor suppressor and an enigma. Genes Dev. 2016 Jul 1;30(13):1492–502. doi: 10.1101/gad.282145.116. PMID: 27401552; PMCID: PMC4949322.

17. Emsley P, Lohkamp B, Scott WG, Cowtan K. Features and development of Coot. Acta Crystallogr D Biol Crystallogr. 2010 Apr;66(Pt 4):486–501. doi: 10.1107/S0907444910007493. Epub 2010 Mar 24. PMID: 20383002; PMCID: PMC2852313.

18. Evans PR. An introduction to data reduction: space-group determination, scaling and intensity statistics. Acta Crystallogr D Biol Crystallogr. 2011 Apr;67(Pt 4):282–92. doi: 10.1107/S090744491003982X. Epub 2011 Mar 18. PMID: 21460446; PMCID: PMC3069743.

19. Fallatah MMJ, Law FV, Chow WA, Kaiser P. Small-molecule correctors and stabilizers to target p53. Trends Pharmacol Sci. 2023 May;44(5):274–289. doi: 10.1016/j.tips.2023.02.007. Epub 2023 Mar 22. PMID: 36964053; PMCID: PMC10511064.

20. Felsani A, Mileo AM, Paggi MG. Retinoblastoma family proteins as key targets of the small DNA virus oncoproteins. Oncogene. 2006 Aug 28;25(38):5277–85. doi: 10.1038/sj.onc.1209621. PMID: 16936748.

21. Henley SA, Francis SM, Demone J, Ainsworth P, Dick FA. A cancer derived mutation in the retinoblastoma gene with a distinct defect for LXCXE dependent interactions. Cancer Cell Int. 2010 Mar 18;10:8. doi: 10.1186/1475-2867-10-8. PMID: 20298605; PMCID: PMC2859746.

22. Huber HE, DeFeo-Jones D, Vuocolo G, Goodhart PJ, Maigetter RZ, Sanyal G, Oliff A, Heimbrook DC. Purification and characterization of a functionally homogeneous 60-kDa species of the retinoblastoma gene product. Journal of Biological Chemistry, Volume 267, Issue 12, 7971–7974. 1992

23. Huynh K, Partch CL. Analysis of protein stability and ligand interactions by thermal shift assay. Curr Protoc Protein Sci. 2015 Feb 2;79:28.9.1-28.9.14. doi: 10.1002/0471140864.ps2809s79. PMID: 25640896; PMCID: PMC4332540.

24. Jarmoskaite I, AlSadhan I, Vaidyanathan PP, Herschlag D. How to measure and evaluate binding affinities. Elife. 2020 Aug 6;9:e57264. doi: 10.7554/eLife.57264. PMID: 32758356; PMCID: PMC7452723.

25. Kabsch W. XDS. Acta Crystallogr D Biol Crystallogr. 2010 Feb;66(Pt 2):125–32. doi: 10.1107/S0907444909047337. Epub 2010 Jan 22. PMID: 20124692; PMCID: PMC2815665.

26. Lee C, Chang JH, Lee HS, Cho Y. Structural basis for the recognition of the E2F transactivation domain by the retinoblastoma tumor suppressor. Genes Dev. 2002 Dec 15;16(24):3199–212. doi: 10.1101/gad.1046102. PMID: 12502741; PMCID: PMC187509.

27. Lee JO, Russo AA, Pavletich NP. Structure of the retinoblastoma tumour-suppressor pocket domain bound to a peptide from HPV E7. Nature. 1998 Feb 26;391(6670):859-65. doi: 10.1038/36038. PMID: 9495340.

28. Liebschner D, Afonine PV, Baker ML, Bunkóczi G, Chen VB, Croll TI, Hintze B, Hung LW, Jain S, McCoy AJ, Moriarty NW, Oeffner RD, Poon BK, Prisant MG, Read RJ, Richardson JS, Richardson DC, Sammito MD, Sobolev OV, Stockwell DH, Terwilliger TC, Urzhumtsev AG, Videau LL, Williams CJ, Adams PD. Macromolecular structure determination using X-rays, neutrons and electrons: recent developments in Phenix. Acta Crystallogr D Struct Biol. 2019 Oct 1;75(Pt 10):861–877. doi: 10.1107/S2059798319011471. Epub 2019 Oct 2. PMID: 31588918; PMCID: PMC6778852.

29. Liu H, Naismith JH. An efficient one-step site-directed deletion, insertion, single and multiple-site plasmid mutagenesis protocol. BMC Biotechnol. 2008 Dec 4;8:91. doi: 10.1186/1472-6750-8-91. PMID: 19055817; PMCID: PMC2629768.

30. McCoy AJ, Grosse-Kunstleve RW, Adams PD, Winn MD, Storoni LC, Read RJ. Phaser crystallographic software. J Appl Crystallogr. 2007 Aug 1;40(Pt 4):658-674. doi: 10.1107/S0021889807021206. Epub 2007 Jul 13. PMID: 19461840; PMCID: PMC2483472.

31. Otterson GA, Chen Wd, Coxon AB, Khleif SN, Kaye FJ. Incomplete penetrance of familial retinoblastoma linked to germ-line mutations that result in partial loss of RB function. Proc Natl Acad Sci U S A. 1997 Oct 28;94(22):12036–40. doi: 10.1073/pnas.94.22.12036. PMID: 9342358; PMCID: PMC23695.

32. Otterson GA, Modi S, Nguyen K, Coxon AB, Kaye FJ. Temperature-sensitive RB mutations linked to incomplete penetrance of familial retinoblastoma in 12 families. Am J Hum Genet. 1999 Oct;65(4):1040–6. doi: 10.1086/302581. PMID: 10486322; PMCID: PMC1288236.

33. Putta S, Alvarez L, Lüdtke S, Sehr P, Müller GA, Fernandez SM, Tripathi S, Lewis J, Gibson TJ, Chemes LB, Rubin SM. Structural basis for tunable affinity and specificity of LxCxE-dependent protein interactions with the retinoblastoma protein family. Structure. 2022 Sep 1;30(9):1340–1353.e3. doi: 10.1016/j.str.2022.05.019. Epub 2022 Jun 17. PMID: 35716663; PMCID: PMC9444907.

34. Pye CR, Bray WM, Brown ER, Burke JR, Lokey RS, Rubin SM. A Strategy for Direct Chemical Activation of the Retinoblastoma Protein. ACS Chem Biol. 2016 May 20;11(5):1192–7. doi: 10.1021/acschembio.6b00011. Epub 2016 Feb 11. PMID: 26845289; PMCID: PMC5117131.

35. Rubin SM. Deciphering the retinoblastoma protein phosphorylation code. Trends Biochem Sci. 2013 Jan;38(1):12–9. doi: 10.1016/j.tibs.2012.10.007. Epub 2012 Dec 3. PMID: 23218751; PMCID: PMC3529988.

36. Rubin SM, Sage J, Skotheim JM. Integrating Old and New Paradigms of G1/S Control. Mol Cell. 2020 Oct 15;80(2):183–192. doi: 10.1016/j.molcel.2020.08.020. Epub 2020 Sep 17. PMID: 32946743; PMCID: PMC7582788.

37. Sagi M, Frenkel A, Eilat A, Weinberg N, Frenkel S, Pe’er J, Abeliovich D, Lerer I. Genetic screening in patients with Retinoblastoma in Israel. Fam Cancer. 2015 Sep;14(3):471–80. doi: 10.1007/s10689-015-9794-z. PMID: 25754945.

38. Sellers WR, Novitch BG, Miyake S, Heith A, Otterson GA, Kaye FJ, Lassar AB, Kaelin WG Jr. Stable binding to E2F is not required for the retinoblastoma protein to activate transcription, promote differentiation, and suppress tumor cell growth. Genes Dev. 1998 Jan 1;12(1):95–106. doi: 10.1101/gad.12.1.95. PMID: 9420334; PMCID: PMC316399.

39. Sherr CJ, Beach D, Shapiro GI. Targeting CDK4 and CDK6: From Discovery to Therapy. Cancer Discov. 2016 Apr;6(4):353–67. doi: 10.1158/2159-8290.CD-15-0894. Epub 2015 Dec 11. PMID: 26658964; PMCID: PMC4821753.

40. Shan B, Durfee T, Lee WH. Disruption of RB/E2F-1 interaction by single point mutations in E2F-1 enhances S-phase entry and apoptosis. Proc Natl Acad Sci U S A. 1996 Jan 23;93(2):679–84. doi: 10.1073/pnas.93.2.679. PMID: 8570615; PMCID: PMC40112.

41. Singh M, Krajewski M, Mikolajka A, Holak TA. Molecular determinants for the complex formation between the retinoblastoma protein and LXCXE sequences. J Biol Chem. 2005 Nov 11;280(45):37868–76. doi: 10.1074/jbc.M504877200. Epub 2005 Aug 23. PMID: 16118215.

42. Sun H, Wang Y, Chinnam M, Zhang X, Hayward SW, Foster BA, Nikitin AY, Wills M, Goodrich DW. E2f binding-deficient Rb1 protein suppresses prostate tumor progression in vivo. Proc Natl Acad Sci U S A. 2011 Jan 11;108(2):704–9. doi: 10.1073/pnas.1015027108. Epub 2010 Dec 27. PMID: 21187395; PMCID: PMC3021049.

43. Sun H, Chang Y, Schweers B, Dyer MA, Zhang X, Hayward SW, Goodrich DW. An E2F binding-deficient Rb1 protein partially rescues developmental defects associated with Rb1 nullizygosity. Mol Cell Biol. 2006 Feb;26(4):1527–37. doi: 10.1128/MCB.26.4.1527-1537.2006. PMID: 16449662; PMCID: PMC1367194.

44. Talluri S, Francis SM, Dick FA. Mutation of the LXCXE binding cleft of pRb facilitates transformation by ras in vitro but does not promote tumorigenesis in vivo. PLoS One. 2013 Aug 6;8(8):e72236. doi: 10.1371/journal.pone.0072236. PMID: 23936539; PMCID: PMC3735560.

45. Talluri S, Isaac CE, Ahmad M, Henley SA, Francis SM, Martens AL, Bremner R, Dick FA. A G1 checkpoint mediated by the retinoblastoma protein that is dispensable in terminal differentiation but essential for senescence. Mol Cell Biol. 2010 Feb;30(4):948–60. doi: 10.1128/MCB.01168-09. Epub 2009 Dec 14. PMID: 20008551; PMCID: PMC2815577.

46. Tate JG, Bamford S, Jubb HC, Sondka Z, Beare DM, Bindal N, Boutselakis H, Cole CG, Creatore C, Dawson E, Fish P, Harsha B, Hathaway C, Jupe SC, Kok CY, Noble K, Ponting L, Ramshaw CC, Rye CE, Speedy HE, Stefancsik R, Thompson SL, Wang S, Ward S, Campbell PJ, Forbes SA. COSMIC: the Catalogue Of Somatic Mutations In Cancer. Nucleic Acids Res. 2019 Jan 8;47(D1):D941–D947. doi: 10.1093/nar/gky1015. PMID: 30371878; PMCID: PMC6323903.

47. Thwaites MJ, Cecchini MJ, Talluri S, Passos DT, Carnevale J, Dick FA. Multiple molecular interactions redundantly contribute to RB-mediated cell cycle control. Cell Div. 2017 Mar 14;12:3. doi: 10.1186/s13008-017-0029-6. PMID: 28293272; PMCID: PMC5348811.

48. Tomita T, Huibregtse JM, Matouschek A. A masked initiation region in retinoblastoma protein regulates its proteasomal degradation. Nat Commun. 2020 Apr 24;11(1):2019. doi: 10.1038/s41467-020-16003-3. PMID: 32332747; PMCID: PMC7181824.

49. Topacio BR, Zatulovskiy E, Cristea S, Xie S, Tambo CS, Rubin SM, Sage J, Kõivomägi M, Skotheim JM. Cyclin D-Cdk4,6 Drives Cell-Cycle Progression via the Retinoblastoma Protein’s C-Terminal Helix. Mol Cell. 2019 May 16;74(4):758–770.e4. doi: 10.1016/j.molcel.2019.03.020. Epub 2019 Apr 11. PMID: 30982746; PMCID: PMC6800134.

50. Vormer TL, Wojciechowicz K, Dekker M, de Vries S, van der Wal A, Delzenne-Goette E, Naik SH, Song JY, Dannenberg JH, Hansen JB, Te Riele H. RB family tumor suppressor activity may not relate to active silencing of E2F target genes. Cancer Res. 2014 Sep 15;74(18):5266–76. doi: 10.1158/0008-5472.CAN-13-3706. Epub 2014 Jul 23. PMID: 25056122.

51. Wallentine BD, Wang Y, Tretyachenko-Ladokhina V, Tan M, Senear DF, Luecke H. Structures of oncogenic, suppressor and rescued p53 core-domain variants: mechanisms of mutant p53 rescue. Acta Crystallogr D Biol Crystallogr. 2013 Oct;69(Pt 10):2146–56. doi: 10.1107/S0907444913020830. Epub 2013 Sep 20. PMID: 24100332; PMCID: PMC3792646.

52. Wan WY, Milner-White EJ. A recurring two-hydrogen-bond motif incorporating a serine or threonine residue is found both at alpha-helical N termini and in other situations. J Mol Biol. 1999 Mar 12;286(5):1651–62. doi: 10.1006/jmbi.1999.2551. PMID: 10064721.

53. Xiao B, Spencer J, Clements A, Ali-Khan N, Mittnacht S, Broceño C, Burghammer M, Perrakis A, Marmorstein R, Gamblin SJ. Crystal structure of the retinoblastoma tumor suppressor protein bound to E2F and the molecular basis of its regulation. Proc Natl Acad Sci U S A. 2003 Mar 4;100(5):2363–8. doi: 10.1073/pnas.0436813100. Epub 2003 Feb 21. PMID: 12598654; PMCID: PMC151346.

54. Zhu Y, Wang A, Allard GM, Nordberg JJ, Nair RV, Kunder CA, Lowe AC. Immunofluorescent and molecular characterization of effusion tumor cells reveal cancer site-of-origin and disease-driving mutations. Cancer Cytopathol. 2022 Oct;130(10):771–782. doi: 10.1002/cncy.22610. Epub 2022 Jun 22. PMID: 35731106.

