## Supplemental Information for "Structural and functional analysis of cancer-associated missense variants in the retinoblastoma protein (Rb) pocket domain" (from another version of this preprint: version 1, 2024-09-22)

### Contents

Supplemental Figure 1

FP binding data for all variants tested — Pg. 1-5

Supplemental Table 1

Primers used in this study — Pg. 6-8

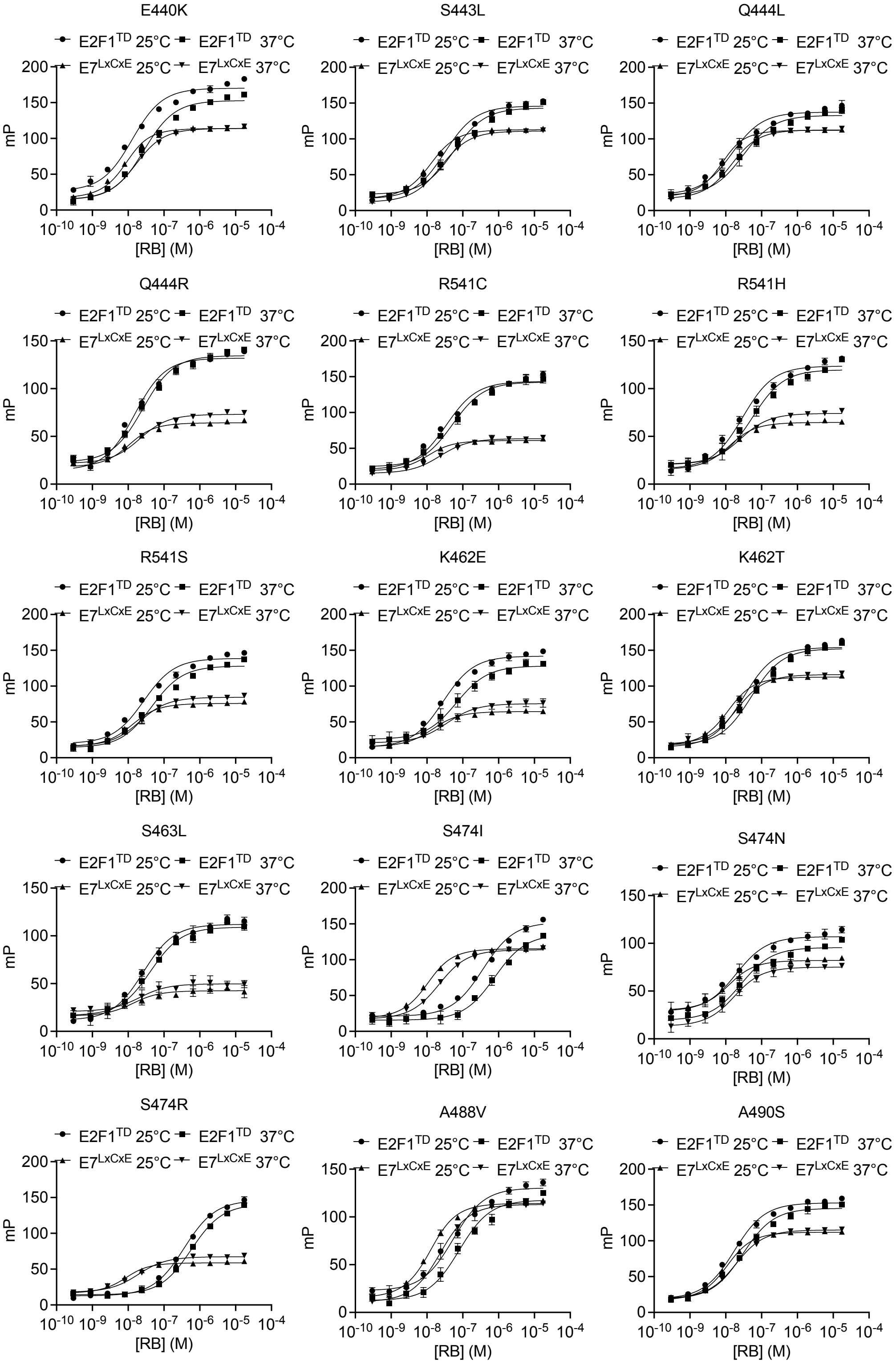


**Supp. Figure 1.** FP data for all variants. Error bars are standard deviations of 4 replicates.

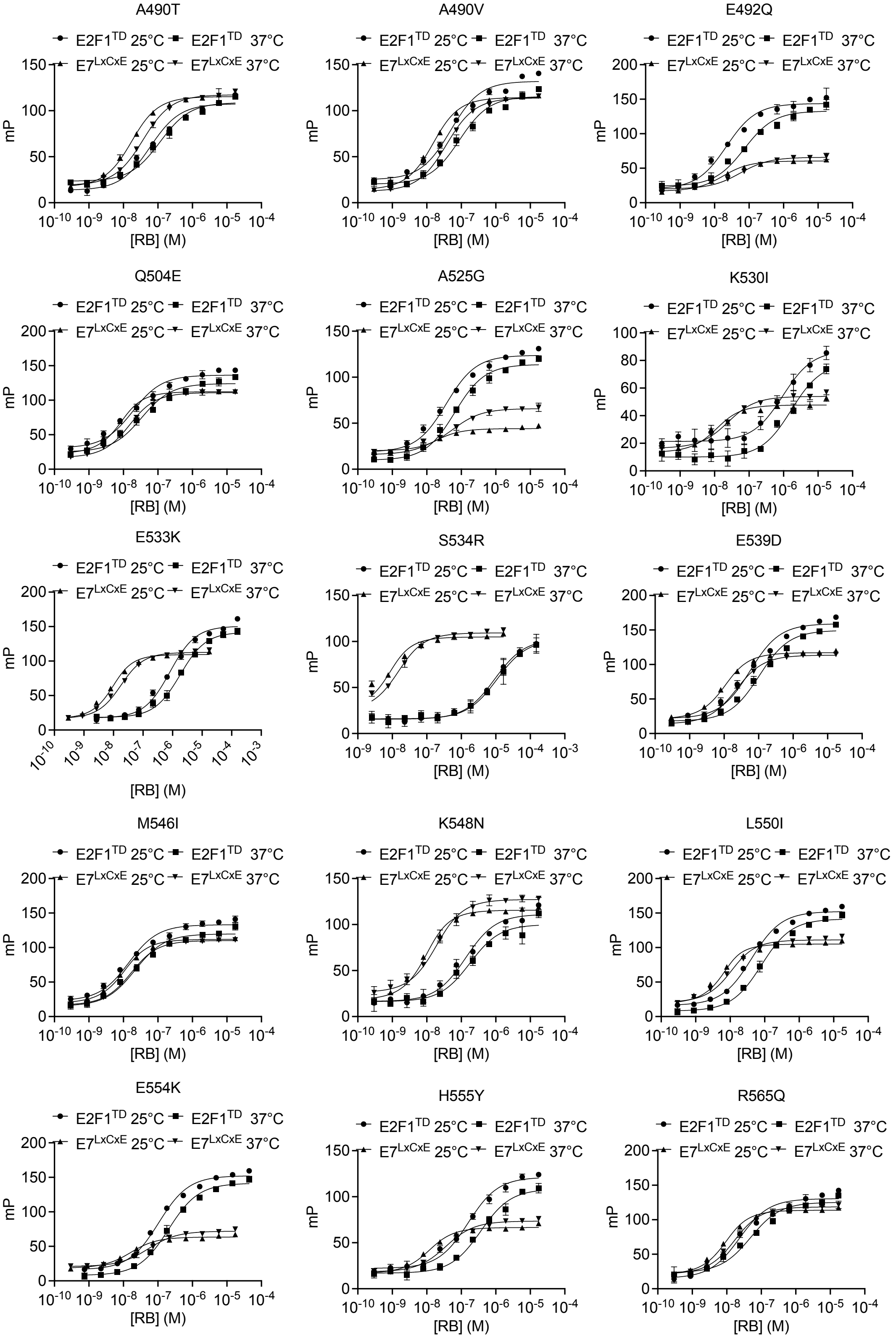


**Supp. Figure 1** (continued)

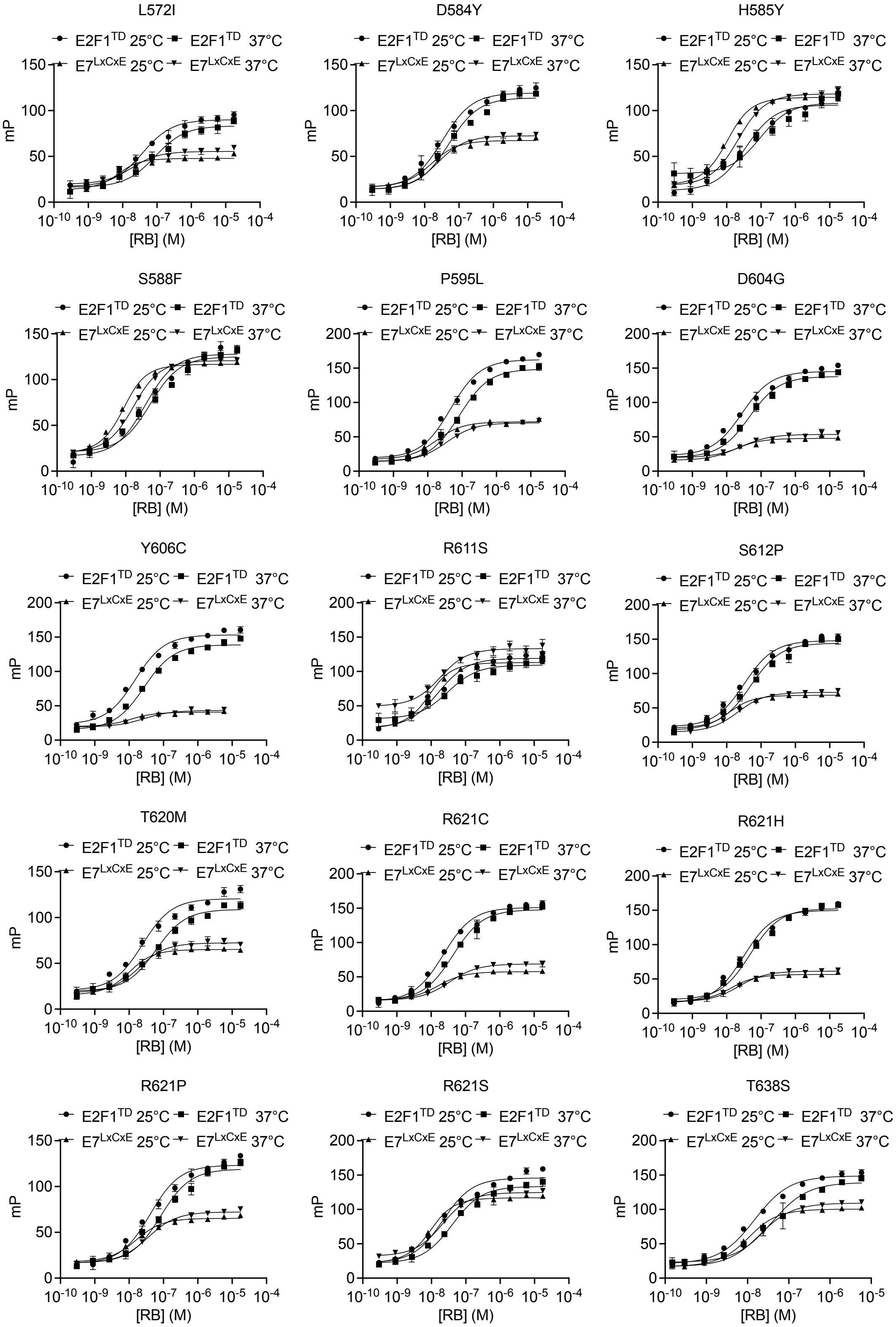


**Supp. Figure 1** (continued)

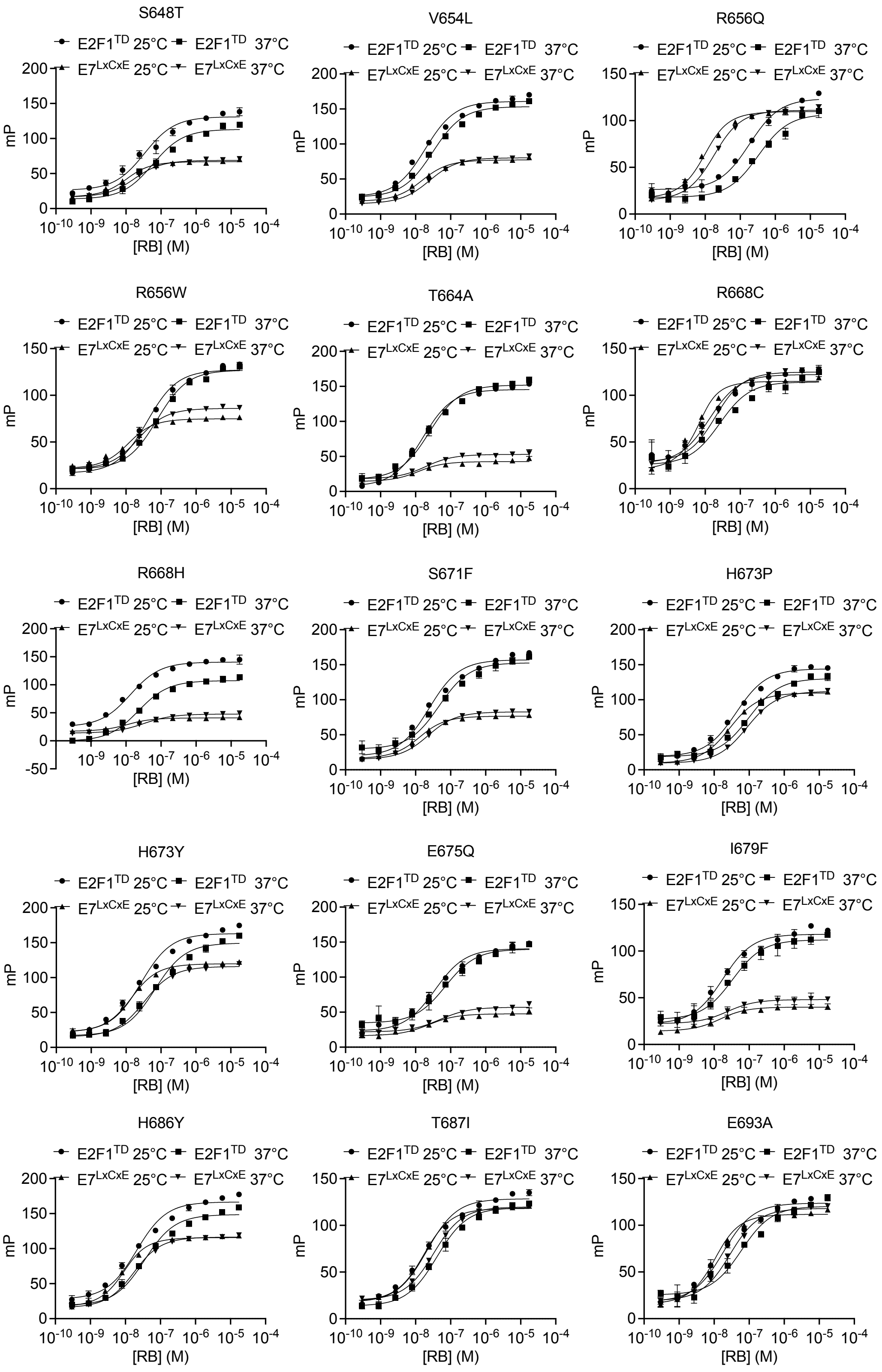


**Supp. Figure 1** (continued)

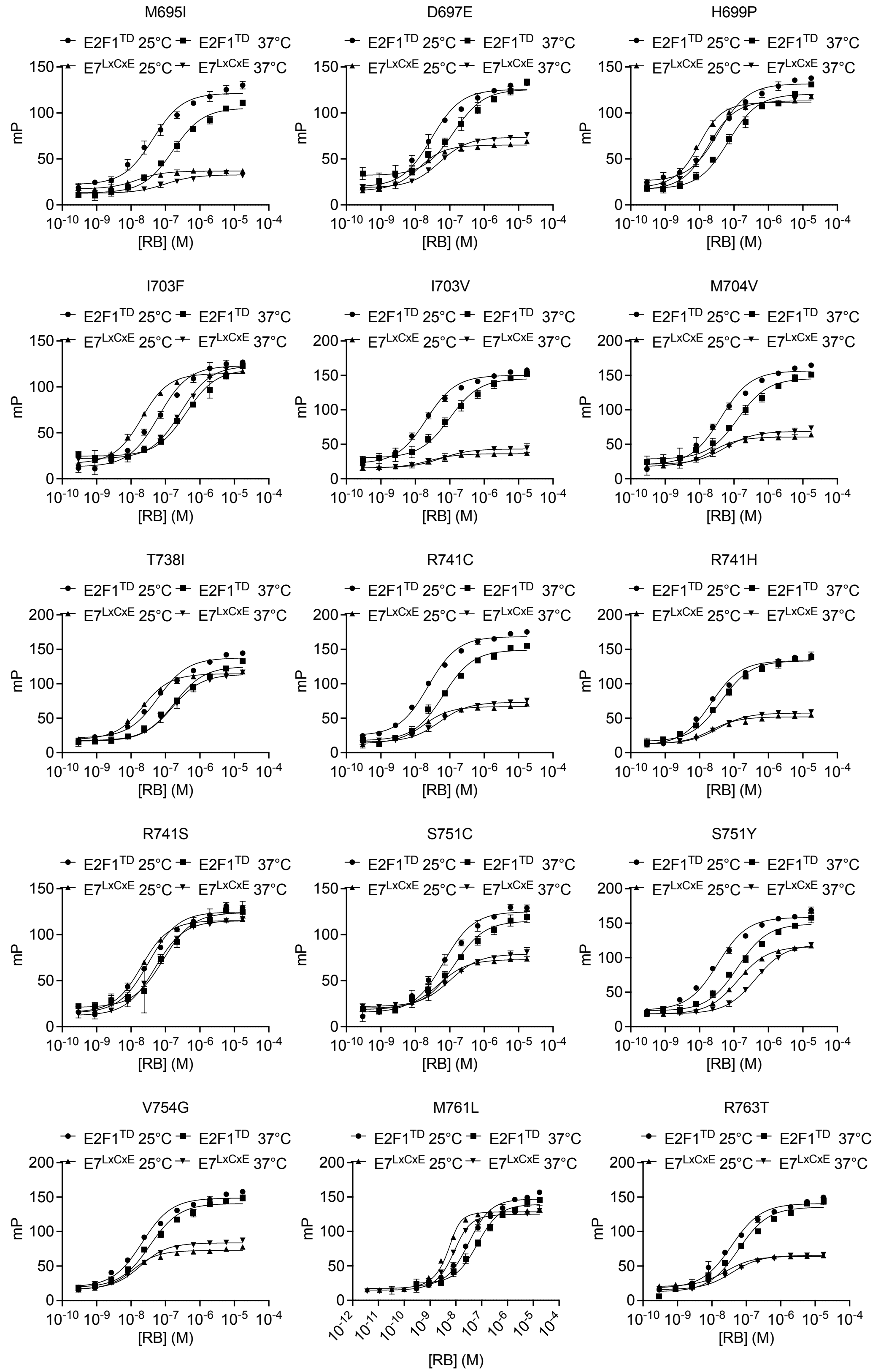


**Supp. Figure 1** (continued)

**Supp. Table 1**. Primers used in this study

| MUTATION | FORWARD PRIMER | REVERSE PRIMER |
|---|---|---|
| E440K | gttgtgtcaaaattggatcacagcgatacaaac | gctgtgatccaattttgacacaaccctgtcc |
| S443L | gaaattggattacagcgatacaaacttgg | cgctgtaatccaatttcgacacaaccctg |
| Q444L | ggatcactgcgatacaaacttggag | gtatcgcagtgatccaatttcgac |
| Q444R | ggatcacggcgatacaaacttggag | gtatcgccgtgatccaatttcgac |
| G449E | gatacaaacttgaagttcgcttgtattaccg | caagcgaacttcaagtttgtatcgctgtgatcc |
| R451C | cttggagtttgcttgtattaccgagtaatgg | cggtaatacaagcaaactccaagtttgtatcgc |
| R451H | cttggagttcacttgtattaccgagtaatgg | cggtaatacaagtgaactccaagtttgtatcgc |
| R451S | cttggagttagcttgtattaccgagtaatgg | cggtaatacaagctaactccaagtttgtatcgc |
| Y454C | cgcttgtattgccgagtaatggaatccatgc | ccattactcggcaatacaagcgaactcc |
| Y454H | cgcttgtatcaccgagtaatggaatccatgc | ccattactcggtgatacaagcgaactcc |
| M457R | cgagtaagggaatccatgcttaaatcag | catggattcccttactcggtaatacaagc |
| L461P | ggaatccatgcctaaatcagaagaagaacg | cttctgatttaggcatggattccattactcgg |
| K462E | ggaatccatgcttgaatcagaagaagaacg | cttctgattcaagcatggattccattactcgg |
| K462T | ggaatccatgcttacatcagaagaagaacg | cttctgatgtaagcatggattccattactcgg |
| S463L | ccatgcttaaattagaagaagaacgattatcc | cttcttctaatttaagcatggattccattac |
| S474I | cattcaaaattttatcaaacttctgaatgacaacatttttc | cagaagtttgataaaattttgaatggataatcgttc |
| S474N | ccattcaaaattttaacaaacttctgaatgacaacatttttc | cagaagtttgttaaaattttgaatggataatcgttc |
| S474R | ccattcaaaattttagaaaacttctgaatgacaacatttttc | cagaagttttctaaaattttgaatggataatcgttc |
| H483R | caacatttttcgtatgtctttattggcgtgcgctcttg | ccaataaagacatacgaaaaatgttgtcattcagaag |
| H483Y | caacatttttatatgtctttattggcgtgcgctcttg | ccaataaagacatataaaaaatgttgtcattcagaag |
| A488E | ctttattggagtgcgctcttgaggttgtaatggcc | caagagcgcactccaataaagacatatg |
| A488V | ctttattggtgtgcgctcttgaggttgtaatggcc | caagagcacacaccaataaagacatatg |
| A490S | ggcgtgctctcttgaggttgtaatggccac | caacctcaagagagcacgccaataaagacatatg |
| A490T | ggcgtgcactcttgaggttgtaatggccac | caacctcaagagtgcacgccaataaagacatatg |
| A490V | ggcgtgcgttcttgaggttgtaatggccac | caacctcaagaacgcacgccaataaagacatatg |
| L491P | cgtgcgctcctgaggttgtaatggccacatatagcag | cattacaacctcaggagcgcacgccaataaagac |
| E492K | cgtgcgctcttaaggttgtaatggccacatatagc | ccattacaaccttaagagcgcacgccaataaagac |
| E492Q | cgtgcgctcttcaggttgtaatggccacatatagc | ccattacaacctgaagagcgcacgccaataaagac |
| Q504E | gtacatctgagaatcttgattctggaacagatttg | gaatcaagattctcagatgtacttctgctatatgtg |
| P515S | gatttgtctttctcatggattctgaatgtgc | gaatccatgagaaagacaaatctgttccagaatc |
| L521P | ctgaatgtgcctaatttaaaagcctttgatttttac | cttttaaattaggcacattcagaatccatggg |
| A525G | cttaatttaaaaggctttgatttttacaaagtgatcg | gtaaaaatcaaagccttttaaattaagcacattcag |
| K530I | gatttttacatagtgatcgaaagttttatcaaagc | cgatcactatgtaaaaatcaaaggcttttaaattaag |
| I532N | caaagtgaacgaaagttttatcaaagcagaag | ctttcgttcactttgtaaaaatcaaag |
| E533K | gtgatcaaaagttttatcaaagcagaaggcaac | gataaaacttttgatcactttgtaaaaatcaaag |
| S534R | gtgatcgaaagatttatcaaagcagaaggcaac | gataaatctttcgatcactttgtaaaaatcaaag |
| E539D | caaagcagacggcaacttgacaagagaaatg | gttgccgtctgctttgataaaactttcg |

| | | |
|---|---|---|
| M546I | caagagaaattataaaacatttagaacgatgtgaacatcg | ctaaatgttttataatttctcttgtcaagttgccttctg |
| K548N | gagaaatgataaaccatttagaacgatgtgaacatcg | gttctaaatggtttatcatttctcttgtcaagttgc |
| L550I | gataaaacatatagaacgatgtgaacatcgaatcatgg | catcgttctatatgttttatcatttctcttgtcaag |
| C553Y | gaacgatatgaacatcgaatcatggaatcccttg | gattcgatgttcatatcgttctaaatgttttatc |
| E554K | gaacgatgtaaacatcgaatcatggaatcccttg | gattcgatgtttacatcgttctaaatgttttatc |
| H555Y | gatgtgaatatcgaatcatggaatcccttgcatg | catgattcgatattcacatcgttctaaatg |
| R556Q | gtgaacatcaaatcatggaatcccttgcatg | catgatttgatgttcacatcgttctaaatg |
| A562E | gaatcccttgaatggctctcagattcacctttatttg | ctgagagccattcaagggattccatgattcgatg |
| A562P | gaatcccttccatggctctcagattcacctttatttg | ctgagagccatggaagggattccatgattcgatg |
| S567L | ctctcagatttacctttatttgatcttattaaac | caaataaaggtaaatctgagagccatgcaaggg |
| L572F | ctttatttgattttattaaacaatcaaaggacc | gtttaataaaatcaaataaaggtgaatctg |
| L572I | ctttatttgatattattaaacaatcaaaggacc | gtttaataatatcaaataaaggtgaatctg |
| D584Y | ggaccaacttatcaccttgaatctgcttgtc | caaggtgataagttggtccttctcggtc |
| H585Y | ccaactgattaccttgaatctgcttgtcctc | gattcaaggtaatcagttggtccttctcggtc |
| S588F | caccttgaatttgcttgtcctcttaatcttcc | ggacaagcaaattcaaggtgatcagttggtcc |
| P595L | cttaatcttcttctccagaataatcacactgcag | gattattctggagaagaagattaagaggacaagcag |
| D604G | ctgcagcaggtatgtatctttctcctgtaagatc | gaaagatacatacctgctgcagtgtgattattc |
| Y606C | cagatatgtgtctttctcctgtaagatctccaaag | ggagaaagacacatatctgctgcagtgtgattattc |
| R611S | ctcctgtaagttctccaaagaaaaaaggttc | ctttggagaacttacaggagaaagatacatatc |
| S612P | ctgtaagacctccaaagaaaaaaggttcaac | ctttggaggtcttacaggagaaagatac |
| T620M | gttcaactatgcgtgtaaattctactgcaaatgc | gaatttacacgcatagttgaacctttttctttg |
| R621C | caactacgtgtgtaaattctactgcaaatgcagag | gaatttacacacgtagttgaacctttttctttg |
| R621H | caactacgcatgtaaattctactgcaaatgcagag | gaatttacatgcgtagttgaacctttttctttg |
| R621P | caactacgcctgtaaattctactgcaaatgcagag | gaatttacaggcgtagttgaacctttttctttg |
| R621S | caactacgagtgtaaattctactgcaaatgcagag | gaatttacactcgtagttgaacctttttctttg |
| T638S | ccttccagtcccagaagccattgaaatctacc | ggcttctgggactggaaggctgaggttgcttg |
| L647R | cctctcgttcactgttttataaaaaagtgtatcggc | cagtgaacgagaggtagatttcaatg |
| V654L | gttttataaaaaactgtatcggctagcctatctc | ctagccgatacagtttttataaaacagtgaaagagag |
| R656Q | gtgtatcagctagcctatctccggctaaatac | gataggctagctgatacactttttataaaacag |
| R656W | gtgtattggctagcctatctccggctaaatac | gataggctagccaatacactttttataaaacag |
| R661Q | cctatctccagctaaatacactttgtgaacgc | gtatttagctggagataggctagccgatacac |
| R661W | cctatctctggctaaatacactttgtgaacgc | gtatttagccagagataggctagccgatacac |
| T664A | ggctaaatgcactttgtgaacgccttctgtc | cacaaagtgcatttagccggagataggctag |
| R668C | ctttgtgaatgccttctgtctgagcacccagaattag | cagaaggcattcacaaagtgtatttagccggag |
| R668H | ctttgtgaacaccttctgtctgagcacccagaattag | gacagaaggtgttcacaaagtgtatttagccggag |
| S671F | ccttctgtttgagcacccagaattagaac | ggtgctcaaacagaaggcgttcacaaagtg |
| H673P | gtctgagcccccagaattagaacatatcatctgg | ctaattctgggggctcagacagaaggcgttcacaaag |
| H673Y | ctgtctgagtacccagaattagaacatatcatctgg | ctaattctgggtactcagacagaaggcgttcacaaag |
| E675Q | gcacccacaattagaacatatcatctggaccc | gatatgttctaattgtgggtgctcagacagaaggc |
| I679F | gaattagaacatttcatctggacccttttccagc | ccagatgaaatgttctaattctgggtgctc |

| | | |
|---|---|---|
| I680T | gaacatatcacctggacccttttccagcacac | gggtccaggtgatatgttctaattctgggtg |
| H686Y | cttttccagtacaccctgcagaatgagtatgaactc | gcagggtgtactggaaaagggtccagatgatatg |
| T687I | ccagcacatcctgcagaatgagtatgaactc | ctgcaggatgtgctggaaaagggtccagatg |
| L688P | gcacaccccgcagaatgagtatgaactcatg | ctcattctgcggggtgtgctggaaaagggtccag |
| E693A | gagtatgcactcatgagagacaggcatttggacc | gagtatgcactcatgagagacaggcatttggacc |
| L694P | gagtatgaacccatgagagacaggcatttggacc | gagtatgaacccatgagagacaggcatttggacc |
| M695I | gaactcattagagacaggcatttggacc | cctgtctctaatgagttcatactcattctgc |
| M695K | gaactcaagagagacaggcatttggacc | cctgtctctcttgagttcatactcattctgc |
| D697E | catgagagagaggcatttggaccaaattatg | caaatgcctctctctcatgagttcatactcattc |
| R698S | gagagacagccatttggaccaaattatgatgtg | gtccaaatggctgtctctcatgagttcatactc |
| R698W | gagagactggcatttggaccaaattatgatgtg | gtccaaatgccagtctctcatgagttcatactc |
| H699P | gagacaggcctttggaccaaattatgatgtg | ggtccaaaggcctgtctctcatgagttcatac |
| D701N | ggcatttgaaccaaattatgatgtgttccatg | cataatttggttcaaatgcctgtctctcatgag |
| D701V | ggcatttggtccaaattatgatgtgttccatg | cataatttggaccaaatgcctgtctctcatgag |
| D701Y | ggcatttgtaccaaattatgatgtgttccatg | cataatttggtacaaatgcctgtctctcatgag |
| Q702K | catttggacaaaattatgatgtgttccatgtatgg | catcataattttgtccaaatgcctgtctctcatgag |
| I703F | ggaccaatttatgatgtgttccatgtatggcatatg | cacatcataaattggtccaaatgcctgtctctc |
| I703V | ggaccaagttatgatgtgttccatgtatggcatatg | cacatcataacttggtccaaatgcctgtctctc |
| M704V | gaccaaattgtgatgtgttccatgtatggcatatg | gaacacatcacaatttggtccaaatgcctgtctc |
| C706F | caaattatgatgttttccatgtatggcatatgcaaagtg | catacatggaaaacatcataatttggtccaaatgcctgtc |
| C706Y | caaattatgatgtattccatgtatggcatatgcaaagtg | catacatggaatacatcataatttggtccaaatgcctgtc |
| M708K | gtgttccaagtatggcatatgcaaagtgaagaatatag | catatgccatacttggaacacatcataatttggtc |
| I724N | caaaatcaatgtaacagcatacaaggatc | gctgttacattgattttgaatttaaggtctatattc |
| I724S | caaaatcagtgtaacagcatacaaggatc | gctgttacactgattttgaatttaaggtctatattc |
| T738I | gttcaggagatattcaaacgtgttttgatcaaag | cacgtttgaatatctcctgaacagcatgagg |
| R741C | cattcaaatgtgttttgatcaaagaagaggag | gatcaaaacacatttgaatgtctcctgaacagcatg |
| R741H | cattcaaacatgttttgatcaaagaagaggag | gatcaaaacatgtttgaatgtctcctgaacagcatg |
| R741S | cattcaaaagtgttttgatcaaagaagaggag | gatcaaaacacttttgaatgtctcctgaacagcatg |
| S751C | gtatgattgtattatagtattctataactcggtc | gaatactataatacaatcatactcctcttctttgatc |
| S751Y | gtatgattatattatagtattctataactcggtc | gaatactataatataatcatactcctcttctttgatc |
| V754G | ctattataggattctataactcggtcttcatgc | gttatagaatcctataatagaatcatactcctcttc |
| Y756C | gtattctgtaactcggtcttcatgcagag | ccgagttacagaatactataatagaatcatac |
| S758L | gtattctataacttggtcttcatgcagagactg | catgaagaccaagttatagaatactataatagaatc |
| M761L | gtcttcctgcagagactgaaaacaaatattttg | gtctctgcaggaagaccgagttatagaatac |
| R763T | catgcagacactgaaaacaaatattttgcag | gttttcagtgtctgcatgaagaccgagttatag |
